# PATHNet: A Thermodynamic Path-Integral-Inspired Neural Network for Protein-Ligand Interaction Prediction

**DOI:** 10.64898/2026.10.11.758399

**Authors:** A. S. Ben Geoffrey, Abhishek Singh, Sowmya Kanchan, Nagaraj M Kulkarni

## Abstract

Protein-ligand scoring functions must capture strongly nonlinear, target-dependent binding physics while remaining generalizable beyond their training data, a balance that purely analytic and purely data-driven models each struggle to achieve on their own. We present PATHNet, a physics-informed framework that learns this unknown functional relationship from molecular-mechanics descriptors using a non-monotonic piecewise-linear mapper with a bounded neural-network correction, trained through a variational, least-action-inspired objective and regulated by a renormalization-group-inspired spectral constraint, with model selection performed through multi-start basin optimization. Across seven DUD-E targets, PATHNet achieved the highest mean active inactive/decoy classification accuracy among the methods tested, outperforming three literature baselines representing distinct architectural families. These results support a physics-grounded, variationally constrained approach to neural-network scoring as a practical route toward more interpretable and transferable protein-ligand interaction models.

## 1. Introduction

Accurate estimation of protein-ligand binding energetics and reliable discrimination of favourable binding configurations remain central problems in structure-based drug discovery. Classical and empirical scoring functions obtain physical interpretability by expressing binding through prescribed analytic terms, typically electrostatic, van der Waals, solvation, hydrogen-bonding, hydrophobic, and strain contributions and then combining these terms through fixed or parameterized functional forms [1-6]. These formulations are efficient and physically motivated, but their predefined functional dependence and often additive structure can restrict the ability of the score to represent strongly nonlinear, target-dependent couplings among the underlying physical variables.

Data-driven machine-learning and deep-learning models address the opposite side of this trade-off. Because multilayer neural networks are flexible universal function approximators, they can learn complex nonlinear associations that are difficult to prescribe analytically [7]. In protein-ligand modelling, however, this flexibility can make the learned relationship strongly dependent on the composition of the training data, and performance can deteriorate when chemical space, target families, or pose distributions differ from those represented during training [1,8,9]. The central objective of the present work is therefore to combine the nonlinear learning capacity of neural methods with an explicitly physics-grounded representation of the binding problem.

PATHNet (Physics-aware Action-based Thermodynamic Hybrid Network) is constructed from this principle. Instead of asking a neural network to infer binding directly from unconstrained molecular descriptors, we define the coordinates of the learning problem using physical quantities known to contribute to protein-ligand binding: intermolecular electrostatic and van der Waals interactions, solvent-mediated contributions, and ligand conformational strain [2,3,10-12]. The binding score is not assumed to be a fixed linear or predetermined analytic combination of these terms. Instead, the relationship between the physics-derived variables and the binding outcome is treated as an unknown function to be learned from data. A trainable non-monotonic piecewise-linear mapper learns this unknown functional form, and a bounded neural-network correction is then added to the function learned by the mapper.

To allow such nonlinear learning while retaining a physical constraint on the solution, we formulate training through a variational correspondence. In classical mechanics and thermodynamics, physically admissible states are characterized by stationarity of an action or thermodynamic potential under the appropriate constraints; the principle of stationary action is expressed by δS = 0 [13,14]. In the present framework, the learned binding functional is sought as a stationary solution over the physics-derived feature space, while the trainable parameters are optimized to a stationary point of the supervised objective. The correspondence is therefore imposed at the level of the variational structure: the model searches for a learned function F that is stationary under the chosen physical representation and simultaneously minimizes prediction error.

A second constraint is introduced through renormalization-group-inspired regulation. Renormalization provides a systematic way to control effective couplings as the description of a many-body system is coarse-grained [15,16]. PATHNet applies this idea operationally to the learned functional expansion: the singular-value spectrum of neural weight matrices and the positive Softplus feature-importance amplitudes are kept within controlled logarithmic-width regimes. This construction is designed to prevent collapse toward numerically singular or spectrally degenerate parameter states and to suppress abrupt scale crossovers in which a small number of couplings dominate the learned solution [15-17].

Related protein-ligand scoring approaches include the physics-informed neural model PIGNet, the geometry-aware attention model GAABind, and the machine-learning force-field static-scoring framework taqo [18-20]; physics-guided neural binding free-energy models have also been reported [21]. The distinctive objective here is to learn an otherwise unknown nonlinear binding functional directly from physics-based feature signatures, solve for that function within a least-action-inspired variational framework, and regulate the learned parameter scales using an RG-inspired scheme. The following Methods section defines the physical decomposition, feature construction, learnable functional mapper and neural correction, variational search, and regulator used to realize this framework.

## 2. Methods

### Physical Formulation of the Learnable Binding Functional

The function approximating the protein-ligand binding free energy to be learned by PATHNet, correlating the target endpoint of binding free energy with the physics-based descriptors, is formulated in terms of non-bonded interaction, solvation, and ligand conformational-strain contributions [2,3,10-12]:

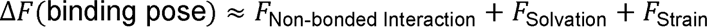

where

- F_Non-bonded Interaction represents intermolecular electrostatic and van der Waals interactions between protein and ligand atoms;
- F_Solvation represents solvent-mediated energetic contributions, including polar and non-polar solvation effects; and
- F_Strain represents the conformational strain energy associated with ligand binding.

The purpose of this decomposition is to define physically meaningful features for learning. The individual energy components and related signatures are therefore supplied to a trainable nonlinear functional approximator. In compact form, the binding relationship is written as

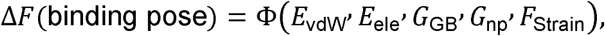

where Φ(·) denotes the unknown nonlinear functional relationship learned during optimization. This formulation allows nonlinear coupling among electrostatic, van der Waals, polar and non-polar solvation, and strain terms without abandoning the physical meaning of the variables from which the function is constructed [2,7,22].

### Variational Solution and Correspondence to the Principle of Least Action

The function learned by PATHNet in approximating protein-ligand binding free energy is physics-grounded in the following sense. In the physical formulation, transition between bound and unbound states may be represented as a variational problem in which the physically admissible trajectory is obtained by stationarity of the corresponding thermodynamic functional. For an equilibrium free-energy functional *F*(α), where α denotes the relevant phase-space coordinates, the stationary condition is

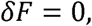

with a stable minimum additionally satisfying

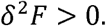

Similarly, in the principle of stationary action, the physical trajectory is selected from the possible paths by requiring

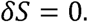

PATHNet is formulated to establish an analogous variational problem in model-parameter space. The learned approximation to the binding functional,

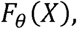

depends on the physics-derived descriptor vector *X* and on the trainable model parameters θ. Different initializations and subsequent variations of θ therefore define different trajectories through the model-parameter space and correspond to different candidate functional solutions.

The parameters are obtained by minimizing the supervised learning objective,

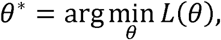

and at convergence the selected solution satisfies the stationarity condition

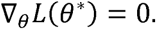

Thus, the correspondence is between two variational-calculus problems. In the physical description, variation of the phase-space coordinates generates alternative admissible physical paths, and the stationary free-energy/action solution is selected. In PATHNet, variation of the model parameters generates alternative learned functional forms, and training selects a stationary solution of the optimization objective. Because the variables supplied to the model are themselves physics-derived quantities describing electrostatic, van der Waals, solvation, and conformational-strain contributions, the learned nonlinear function remains defined over a physically meaningful coordinate system.

The correspondence can be summarized as follows in Table 1 as below

**Table 1.** Correspondence between the neural network and physics based solution.

| Equivalence of two variational-calculus problems | Physics | Neural Network |
| --- | --- | --- |
| Coordinates varied | Phase-space coordinates $\alpha$ | Model parameters $\theta$ |
| Functional representation | Free-energy functional $F(\alpha)$ | Learned functional $F_\theta(X)$ |
| Variational search | Variation over admissible physical paths | Variation over parameter-space trajectories |
| Optimization principle | Action/free-energy minimization | Loss minimization |
| Stationary condition | $\delta F = 0, \delta S = 0$ | $\nabla_\theta L(\theta^*) = 0$ |
| Stable solution | $\delta^2 F > 0$ | Converged low-loss parameter basin |
| Selected solution | Stationary physical trajectory | Converged PATHNet functional |

Because the neural-network objective is non-convex, a single initialization need not identify the most favourable stationary solution. PATHNet therefore begins from multiple independent initializations of θ. Short optimization allows these trajectories to relax toward different local minima of the loss landscape, after which the retained low-loss basins are subjected to extended optimization. The resulting converged solutions are then compared using the validation criterion. In this way, the multi-start optimization procedure provides the computational analogue of exploring multiple admissible variational paths before selecting the most favorable converged solution.

### Dataset Construction and Learning Objective

The learning problem follows an active and decoy dataset construction based on protein-ligand crystal complexes drawn from PDBbind [23]. Complexes with reported ligand potency of 1 µM or better were selected for the starting dataset. For each selected complex, the crystallographic ligand was re-docked into its cognate binding site using AutoDock Vina [24]. A re-docked pose with a heavy-atom root-mean-square deviation (RMSD) of less than 3.0 Å from the crystallographic pose was designated as the active/native-like pose, whereas alternative low-energy Vina poses were retained as decoys (inactives). This produces, for each PDB complex, an active pose together with a set of energetically plausible docking decoys for supervised discrimination. The model therefore learns a bounded score that distinguishes native-like configurations from low-energy decoys using physics-derived features rather than raw coordinates. External validation is performed on seven DUD-E targets [25]: heat shock protein 90-alpha (HSP90A; labeled HSPA in the supplied panels), androgen receptor (AR), dopamine D3 receptor (DRD3), serine/threonine-protein kinase AKT1 (AKT1), cyclin-dependent kinase 2 (CDK2), tyrosine-protein kinase ABL1 (ABL1), and adenosine A2A receptor (AA2AR).

For prepared dataset of actives and decoys in each PDB block of PDBBind, the feature construction is carried out as follows:

### Physics-Based Feature Signatures

For every active or decoy pose, physically interpretable feature signatures are derived from molecular mechanics-based principles. The implementation follows the non-bonded, solvation, and ligand-strain feature construction below [2,3,10-12].

### (a) Non-bonded Interaction Feature Signatures

Molecular-mechanics force-field parameters are first assigned to the protein-ligand complex. The assigned parameters comprise Amber atom types, partial atomic charges, Lennard-Jones parameters, bonded parameters, and molecular topology [3]. The intermolecular non-bonded interaction energy is computed as

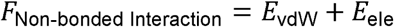

where

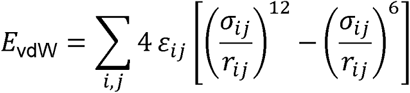

and

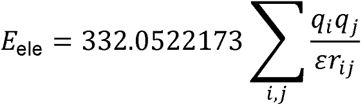

where

- r_ij denotes the distance between protein atom i and ligand atom j;
- q_i and q_j denote the corresponding Amber partial atomic charges;
- ε_ij and σ_ij denote the Lennard-Jones parameters obtained from the assigned molecular-mechanics force field; and
- ε denotes the dielectric constant.

The computed electrostatic and van der Waals interaction energies collectively define the non-bonded interaction feature signature supplied to the learning architecture [2,3].

### (b) Solvation Feature Signatures

To account for solvent effects during molecular recognition, physics-based solvation descriptors are computed using a Generalized Born implicit-solvent model [12]. The total solvation contribution is represented as

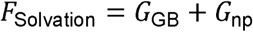

where the polar solvation contribution is calculated as

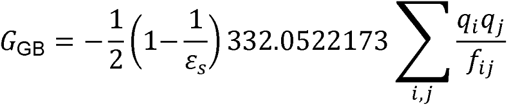

where

- q_i and q_j are Amber partial atomic charges;
- ε_s is the solvent dielectric constant, typically 80 for water; and
- f_ij is the Generalized Born effective distance between atoms.

The non-polar solvation contribution is estimated from solvent-accessible surface area (SASA) [11]:

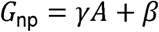

where

- A is the solvent-accessible surface area;
- γ is a surface-tension coefficient (approximately 0.005 kcal mol^-1 Å^-2); and
- β is an empirical constant.

The polar and non-polar terms collectively define the solvation feature signature used by PATHNet [11,12].

### (c) Ligand Conformational Strain Feature Signatures

The conformational-strain contribution quantifies the energetic penalty associated with ligand deformation upon binding. Ligand strain is estimated as the difference between the molecular-mechanics energy of the ligand in its bound conformation and the energy of the corresponding locally minimized conformation [10]:

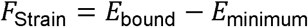

where

- E_bound denotes the molecular-mechanics energy of the ligand in the experimentally observed or docked binding pose; and
- E_minimum denotes the energy of the corresponding locally minimized ligand conformation.

The strain feature therefore represents the energetic cost required for the ligand to adopt the bound geometry within the protein binding pocket.

### Construction of the Physics-Based Feature Vector

For every active and decoy pose, the computed non-bonded interaction, solvation, and strain signatures are assembled into a physics-derived feature vector

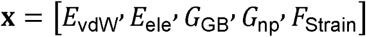

or into higher-dimensional descriptor vectors derived from the same physical quantities. The resulting representation is supplied to the learnable functional mapper, which estimates Φ from these physics-based coordinates rather than from an unconstrained latent representation.

The neural network construction that allows for the learning physics grounded function approximation binding affinity to be able to differentiate actives and in-actives is as follows :

### Hybrid NM-PWL Mapper and Neural-Network Correction Architecture

The central functional approximator is a Non-Monotonic Piecewise Linear (NM-PWL) mapper. Piecewise-linear representations provide flexible approximation of continuous functions on compact domains [7,22]. Here, the segment slopes are unconstrained in sign, so the learned relationship may increase, decrease, or change direction as the effective input varies. The mapper is used to learn the otherwise unknown nonlinear functional relationship between the physics-derived descriptors and the binding score. For normalized scalar input z on a compact interval, a convenient continuous representation is

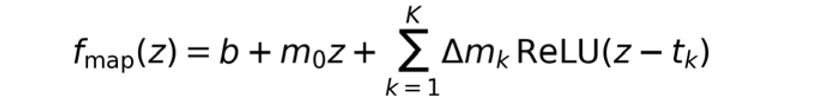

The ordered knots define the breakpoints of the piecewise-linear function, the initial slope defines the response before the first knot, and learned slope increments modify the response after successive knots. Because these increments are not sign-constrained, the mapper can represent increasing, decreasing, convex, concave, multimodal, and non-monotonic relationships while retaining an explicit piecewise-interpretable functional form.

Because the segment slopes are unconstrained, the learned NM-PWL response is not restricted to a single qualitative shape. Depending on the physics-derived input, the mapper can learn increasing, decreasing, convex, concave, multimodal, or non-monotonic relationships. Representative examples from the earlier formulation are shown below: a peaked non-monotonic response with an interior optimum and a monotonic convex response.

**Figure 1A.**
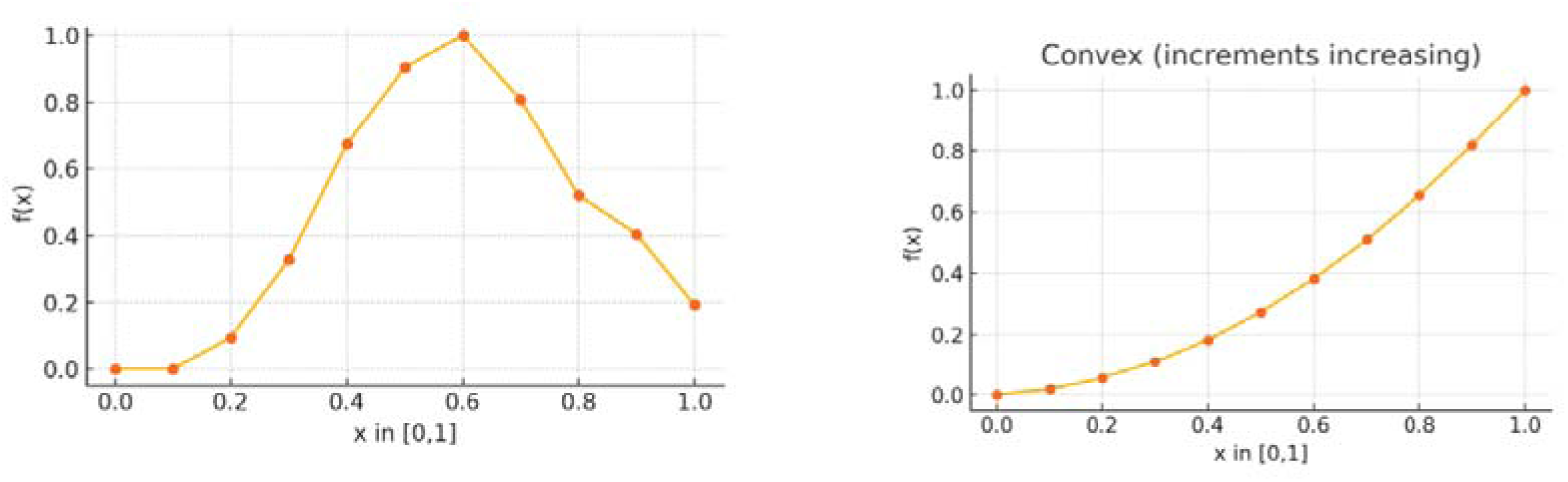
Representative functional forms that may be learned by the NM-PWL mapper. Left: a non-monotonic response with an interior maximum. Right: a monotonic convex response. The learned mapper is not restricted to either profile; its trainable segment parameters adapt the functional form to the physics-derived descriptors.

### Neural-Network Correction Architecture

In addition to the NM-PWL functional mapper, PATHNet contains a separate feed-forward neural-network pathway that provides a flexible data-driven correction to the physics-based representation. The input physics descriptor vector x is first projected into a 32-dimensional latent representation, followed by successive hidden layers of 16 and 4 neurons. Hyperbolic-tangent activation functions are applied after each hidden transformation, giving the architecture

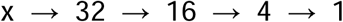

The corresponding transformations are

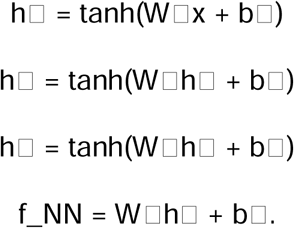

This 32→16→4→1 structure uses tanh nonlinearities after the 32-, 16-, and 4-dimensional layers, followed by a final linear scalar output. The progressively smaller hidden representations provide a hierarchical compression of the input physics descriptors. The first 32-dimensional layer allows the network to construct a comparatively rich latent representation from combinations of the physical variables. The subsequent 16- and 4-dimensional layers progressively compress this representation, retaining combinations of features that are most useful for discriminating native-like and decoy poses.

For active and decoy classification task, the scalar network output is treated as a logit and transformed into a probability through the sigmoid function

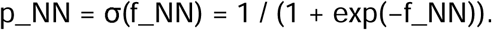

### Learnable Feature Importance Through Softplus Weighting

The relative importance of the input physics-based descriptors is learned directly during network training. Each descriptor is associated with a trainable scalar parameter, which is optimized jointly with the remaining PATHNet parameters through backpropagation. To obtain positive and normalized feature-importance weights, the trainable parameters are transformed using the Softplus function and subsequently normalized:

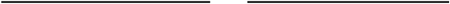

where is the number of input descriptors. By construction,

The resulting weights therefore provide a learned, normalized measure of the relative contribution of each physics-based descriptor to the PATHNet representation.

### Bounded Neural-Network Correction

After the NM-PWL mapper has learned the nonlinear functional relationship from the physics-derived inputs, a secondary feed-forward neural network is used to learn a bounded correction to the mapper-learned function. The two outputs are combined through the convex correction specified in the architecture:

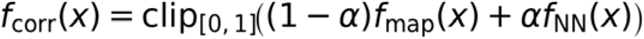

Here, f_map is the function learned by the NM-PWL mapper, f_NN is the bounded neural-network correction, and α is a learned mixing coefficient constrained to an admissible interval. The final prediction therefore retains the explicitly learned piecewise functional relationship while allowing an additional data-driven correction wherever this improves the supervised objective.

**Figure 1B.**
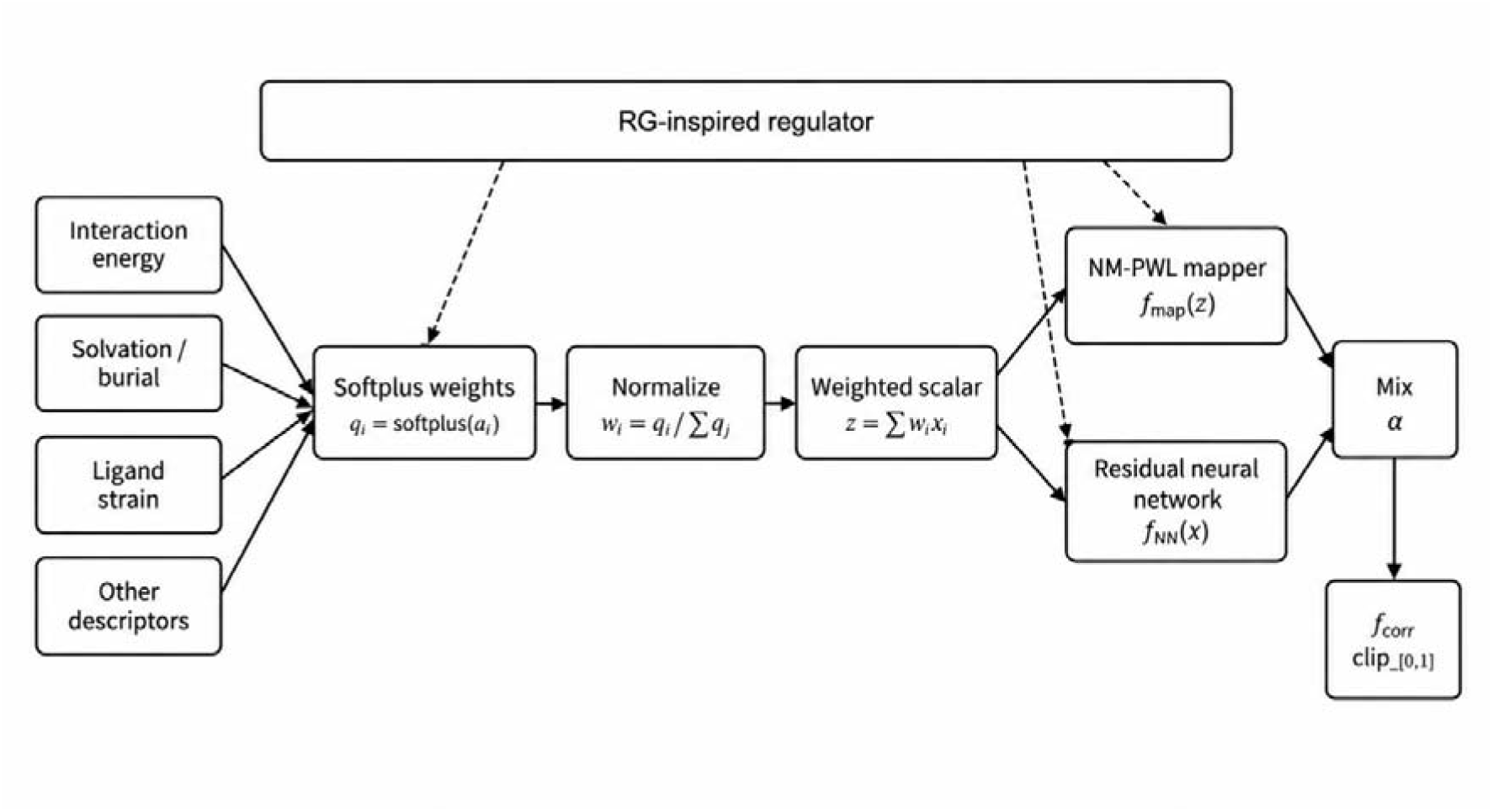
Hybrid NM-PWL + neural-network correction architecture. Physics descriptors are aggregated using normalized Softplus importance weights, mapped by the non-monotonic piecewise-linear functional, and corrected by a bounded residual neural network. The RG-inspired regulator acts on both neural-network weights and the mapper-importance vector.

### Multi-Start Optimization Through Multiple Initializations and Basin Search

Training deep neural networks corresponds to minimizing a highly non-convex loss function

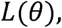

where

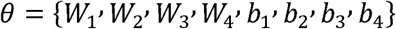

collectively denotes all trainable parameters.

The optimization landscape generally contains numerous local minima,

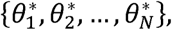

each corresponding to a different approximation of the underlying free-energy functional.

To reduce sensitivity to initialization, PATHNet begins from N independent initializations of the trainable parameter set, θ_i^(0), i = 1, …, N. Each initialization is optimized so that its trajectory relaxes toward a candidate local minimum (attraction basin) of the non-convex loss landscape. The top-k low-loss basins identified during this screening stage are retained and subsequently propagated through extended optimization. The extended stage allows the candidate solutions to converge more fully, after which the final basin is selected using the combined BCE, AUROC, and Top-1 criterion defined below.

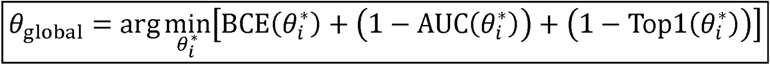

Here, θ_i* denotes the converged parameter set obtained from the i-th retained basin. BCE loss corresponds to Binary Cross Entropy loss corresponding to the i-th retained basin. AUC(θ_i*) is the validation area under the ROC curve for discrimination of active poses from decoy poses across the validation set. Top1(θ_i*) measures pose-ranking success at the PDB-complex level: for each complex containing one active pose and its associated decoys, Top-1 is successful when the active pose receives the highest predicted score among all poses in that complex;

From the perspective of variational optimization, each initialization defines an independent trajectory through parameter space. The initial training stage identifies distinct candidate local minima, and the extended stage allows each retained basin to converge more fully before validation-based model selection.

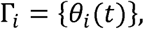

This selection rule is the neural-network analogue of choosing, among the admissible explored trajectories, the stationary solution with the lowest effective action.

### Orthogonal Optimization and Functional Basis Construction

According to functional analysis [26,27] the unknown free-energy functional is represented as

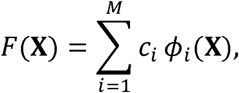

where

- *c_i_* are expansion coefficients,
- *ϕ_i_*(*X*) denote basis functions implicitly learned by the neural network.

Unlike conventional optimization algorithms, the present implementation uses orthogonalized matrix updates for weight matrices, while biases are updated with AdamW [28].

The objective is to construct parameter directions that remain approximately orthogonal throughout optimization,

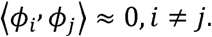

Consequently,

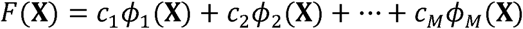

becomes analogous to classical orthogonal expansions used throughout mathematical physics,

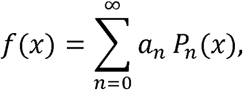

where *P_n_*(*x*) denotes Legendre polynomials,

or

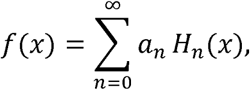

for Hermite polynomials,

or

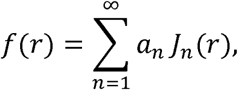

for Bessel function expansions.

Rather than prescribing these basis functions analytically, the neural network learns the basis directly from data while the Muon optimizer encourages decorrelated parameter directions. Consequently, the learned representation behaves as an adaptive orthogonal functional expansion of the unknown binding free-energy surface.

### Renormalization-Based Spectral Weight Regulation

Following every optimization step, a dynamic RG-inspired regulation procedure is applied to the trainable coupling parameters. For neural-network weight matrices, regulation acts on the singular-value spectrum; for the positive Softplus mapper feature-importance amplitudes, an analogous vector log-spread regulator is applied before normalization. Unlike conventional weight decay, which uniformly penalizes parameter magnitudes, this procedure controls relative mode amplitudes. By bounding the logarithmic separation between the largest and smallest effective modes, the regulator is designed to keep the optimization trajectory away from singular-value collapse, spectrally degenerate states, and abrupt scale crossovers in which a small number of couplings dominate the solution [15-17].

For every weight matrix

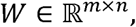

the singular value decomposition is computed,

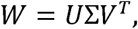

where

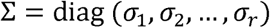

contains the ordered singular values

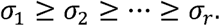

Rather than constraining individual weights, the regulator constrains the **spectral width**

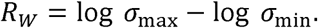

The network is required to remain inside a prescribed spectral operating region,

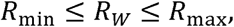

where the present implementation employs

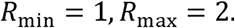

When the spectrum becomes excessively broad,

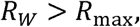

the smallest singular value is increased toward

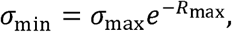

through

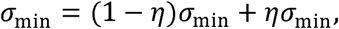

while spectra becoming excessively flat,

are similarly adjusted toward

The regulated weight matrix is reconstructed as

thereby preserving the learned singular vectors while modifying only the effective coupling strengths represented by the singular values.

### Regulation of Softplus Mapper-Importance Weights

The same renormalization-inspired principle is applied to the positive Softplus mapper amplitudes. Because these amplitudes form a vector rather than a matrix, regulation is performed directly in log-amplitude space instead of through a singular-value decomposition. The corresponding log-spread is defined as

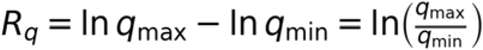

After each optimizer step, the mapper-amplitude log-spread is constrained to the same controlled operating interval used for the neural spectral width. If the vector becomes too broad, extreme amplitudes are multiplicatively compressed toward the admissible boundary; if it becomes too flat, the spread is restored toward the lower boundary. The regulated positive amplitudes are then normalized to unit sum before being supplied to the mapper. This limits single-descriptor domination while avoiding collapse to effectively indistinguishable feature weights.

### Spectral Operating Region and Connection to Random Matrix Theory

Random Matrix Theory and heavy-tailed spectral analyses have shown that the empirical spectral distribution of neural-network weight matrices can provide information about conditioning, scale separation, and effective model complexity [17]. Networks exhibiting excessively sharp spectra may be dominated by a small number of highly amplified modes, whereas overly flat spectra can correspond to under-utilization of representational capacity.

The proposed regulator therefore maintains the learned operators within a controlled spectral operating region by continuously enforcing

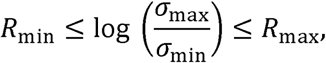

thereby preventing transitions toward either poorly conditioned or spectrally collapsed regimes.

Unlike post hoc WeightWatcher analysis, which estimates quantities such as the Heavy-Tailed power-law exponent

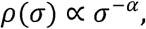

to characterize trained networks, the present method actively regulates the singular-value spectrum throughout optimization. Consequently, rather than merely diagnosing spectral quality after training, the regulator continuously steers the optimization trajectory toward a stable spectral phase consistent with the conditioning principles emphasized by Random Matrix Theory.

### Regulation of Parameter Flow

The regulator additionally constrains the magnitude of parameter updates.

Let

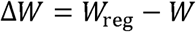

denote the update field.

Its logarithmic spread is computed as

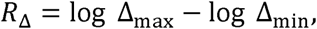

where

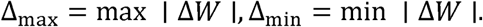

Whenever

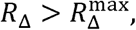

the largest updates are clipped,

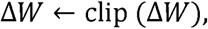

ensuring that optimization proceeds through smooth collective parameter evolution rather than abrupt movements dominated by a small number of weights.

### Renormalization and Effective Theory Interpretation

Within Wilsonian renormalization, microscopic degrees of freedom are progressively coarse-grained into effective collective modes that govern macroscopic observables [15,16]. The effective action is expressed as

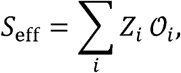

where

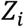

represent renormalized coupling constants associated with the effective operators

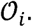

The proposed neural network admits a closely analogous interpretation. The left and right singular vectors,

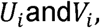

define an orthogonal operator basis learned from the data, while the singular values

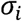

represent the effective coupling strength associated with each learned operator. The learned free-energy functional may therefore be written as

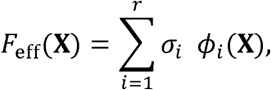

where

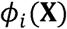

denotes the nonlinear basis generated by the corresponding singular mode.

Dynamic spectral regulation continuously renormalizes these effective couplings by maintaining

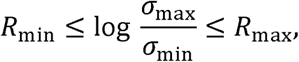

thereby preventing any single microscopic mode from dominating the learned representation while simultaneously preserving the collective modes responsible for the macroscopic approximation of the binding free-energy landscape.

From this perspective, the neural network is interpreted not as a direct microscopic representation of the underlying many-body interaction, but as a renormalized effective theory whose parameters evolve within a controlled spectral phase throughout optimization [15,16].

The matrix component of the regulator is motivated by Random Matrix Theory and heavy-tailed spectral analyses of deep neural networks [17]. Unlike post-training diagnostic tools, the present implementation performs active regulation during optimization. The mapper-weight extension applies the same coarse-graining philosophy to the positive Softplus importance amplitudes, so both residual-network couplings and interpretable mapper couplings are kept inside controlled operating regimes.

### Stepwise Methods Summary

#### 1. Dataset selection: PDBbind

PDBbind complexes with reported ligand potency of 1 µM or better were selected as the starting dataset. The corresponding PDB identifiers define the protein-ligand complexes taken forward for redocking and PATHNet feature generation.

#### 2. Active-decoy dataset generation: AutoDock Vina redocking

The crystallographic ligand was redocked into its cognate PDBbind receptor using AutoDock Vina. Docked poses with heavy-atom RMSD <3 Å from the crystallographic ligand were labeled native-like/active, while other retained low-energy docked clusters were used as decoys/inactives.

#### 3. PATHNet physics-based feature generation

Amber/GAFF-derived parameters were used to calculate physically interpretable descriptors describing electrostatic interaction, van der Waals interaction, solvation, and ligand conformational strain for each protein-ligand pose. These quantities constitute the physics-based input representation supplied to PATHNet.

#### 4. PATHNet modelling

The calculated physics descriptors were supplied to the PATHNet architecture, which combines learned feature weighting, the NM-PWL functional mapper, and a neural-network pathway to learn the nonlinear relationship between the physical descriptors and native-like versus decoy binding poses. An RG-inspired spectral regulator was additionally applied during optimization to control the model weight spectrum and promote convergence toward more stable solutions.

#### 5. External validation: DUD-E

The trained model was externally evaluated on seven DUD-E targets spanning several protein families: HSP90A ( https://dude.docking.org/targets/hs90a), androgen receptor (AR) ( https://dude.docking.org/targets/andr), DRD3 (https://dude.docking.org/targets/drd3), AKT1 ( https://dude.docking.org/targets/akt1), CDK2 ( https://dude.docking.org/targets/cdk2), ABL1 ( https://dude.docking.org/targets/abl1), and AA2AR ( https://dude.docking.org/targets/aa2ar). This set was used to assess whether the learned physics-informed scoring relationship transfers across structurally different binding pockets and target classes.

The codes associated with these steps are shared in the supplementary files.

## 3. Results and Discussion

### 3.1 Effect of RG regulation on loss-basin discovery and extended basin optimization

The multi-start basin-search procedure was designed to reduce dependence on a single random initialization and to test whether RG-inspired regulation reshapes the accessible optimization landscape. The independent parameter initializations were subjected to optimization to identify candidate local minima for subsequent basin comparison. This process was carried out with and without the RG regulator to assess its effect on the accessible optimization landscape. Without the regulator, the identified local minima ranged from 0.67 to 0.57, as shown in Table 2; with the regulator, the corresponding values ranged from 0.63 to 0.43. All retained basins were then subjected to extended optimization; the lowest converged losses obtained without and with the regulator were 0.55 and 0.37, respectively.

**Figure 2.**
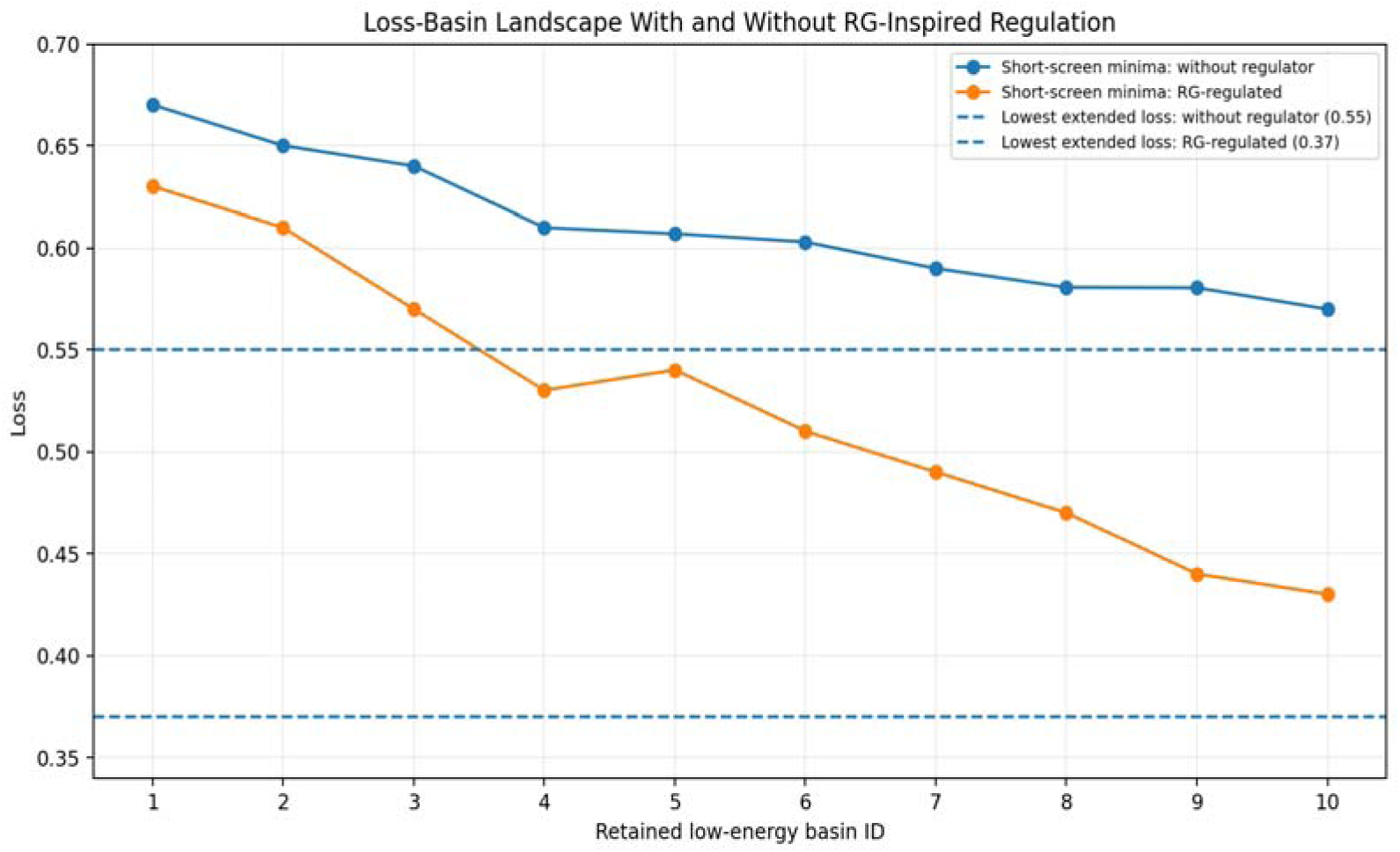
Loss-basin landscape with and without RG-inspired regulation. Solid lines show the short-screen minimum loss for each of the ten retained low-energy basins. Dashed horizontal lines denote the lowest converged losses obtained only after all retained basins were run for extended epochs: 0.55 without the regulator and 0.37 with the regulator. These extended-stage values are ensemble-level outcomes and are not assigned to any single short-screen basin.

**Table 2.**
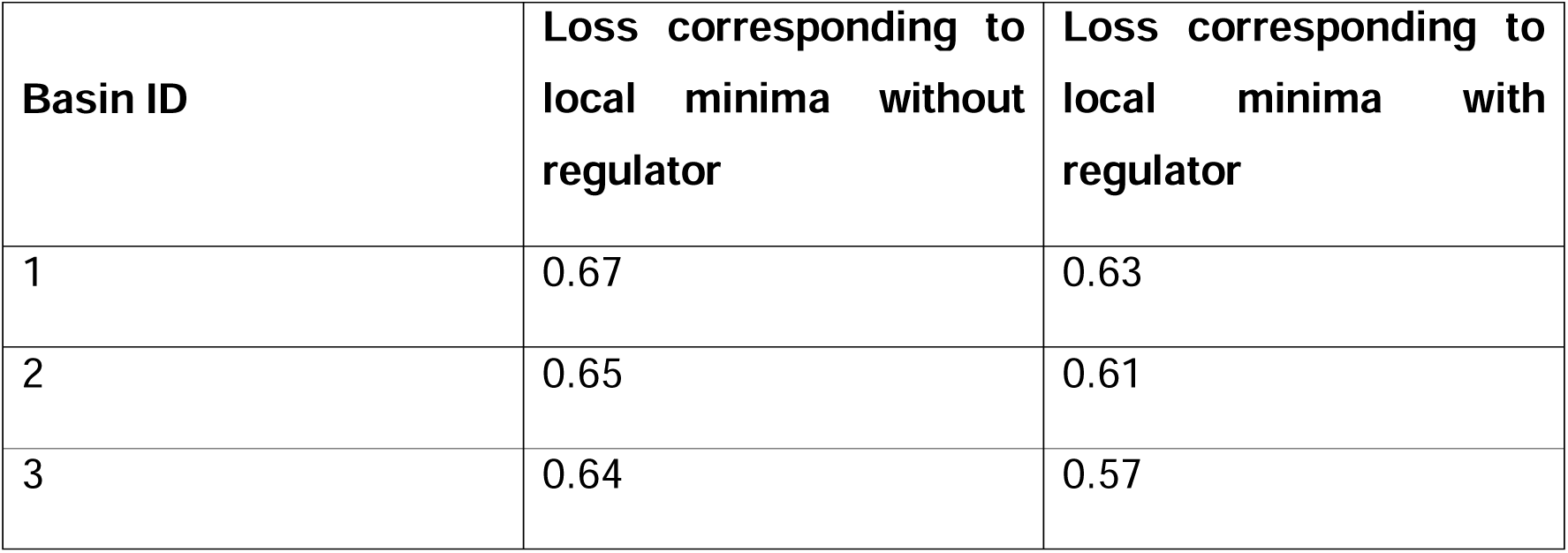
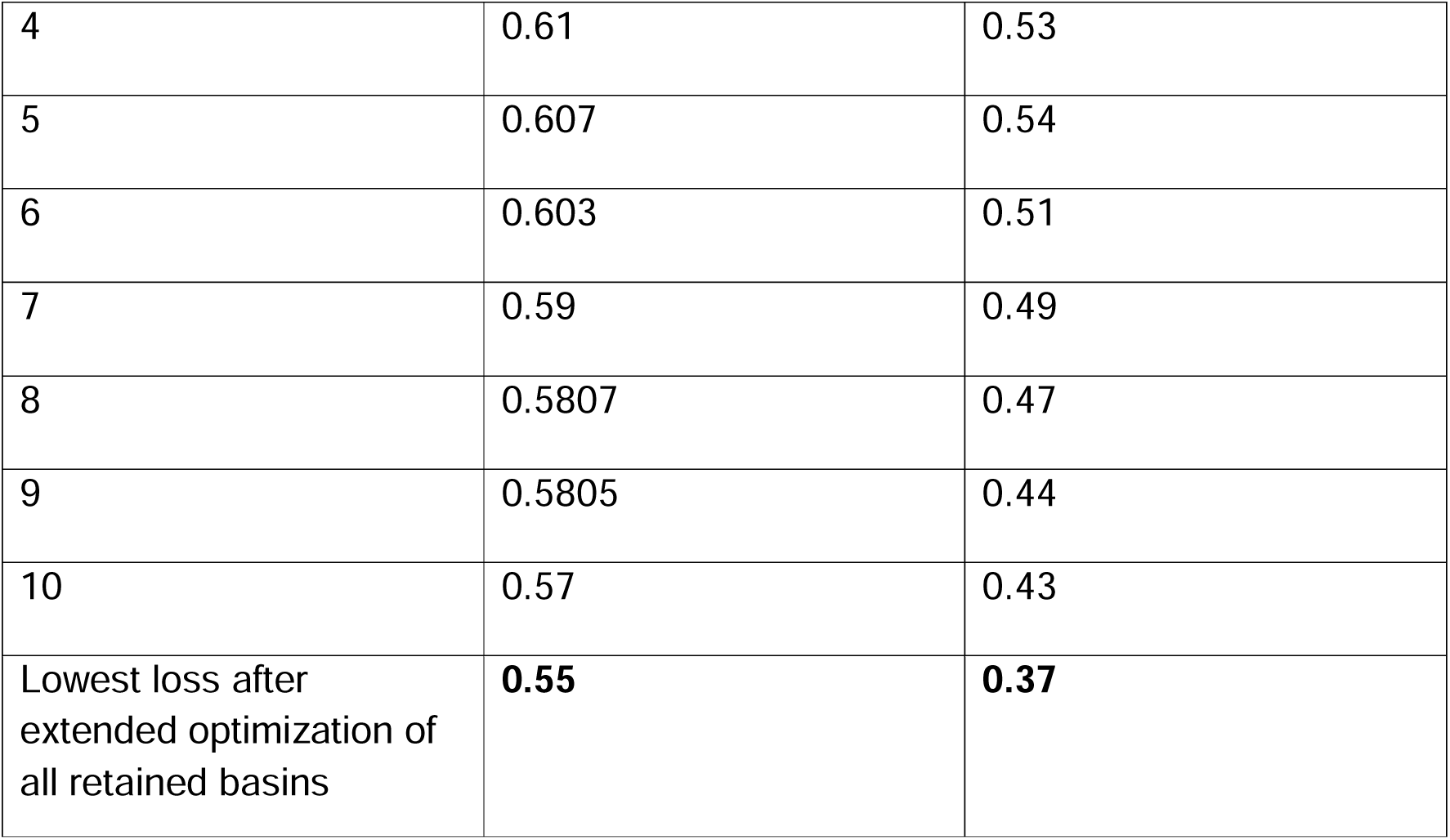
The local minima basins with and without the RG-inspired regulator, together with the lowest converged losses obtained after extended optimization of all retained basins.

| Basin ID | Loss corresponding to local minima without regulator | Loss corresponding to local minima with regulator |
| --- | --- | --- |
| 1 | 0.67 | 0.63 |
| 2 | 0.65 | 0.61 |
| 3 | 0.64 | 0.57 |
| 4 | 0.61 | 0.53 |
| 5 | 0.607 | 0.54 |
| 6 | 0.603 | 0.51 |
| 7 | 0.59 | 0.49 |
| 8 | 0.5807 | 0.47 |
| 9 | 0.5805 | 0.44 |
| 10 | 0.57 | 0.43 |
| Lowest loss after extended optimization of all retained basins | **0.55** | **0.37** |

### 3.2 External validation across seven supplied DUD-E targets

External validation was assessed using seven supplied DUD-E target panels [25]: heat shock protein 90-alpha (HSP90A; labeled HSPA in the supplied panels), androgen receptor (AR), dopamine D3 receptor (DRD3), serine/threonine-protein kinase AKT1 (AKT1), cyclin-dependent kinase 2 (CDK2), tyrosine-protein kinase ABL1 (ABL1), and adenosine A2A receptor (AA2AR). Each panel reports active-versus-inactive score distributions and kernel-density-estimate (KDE)-based thresholded accuracy for PATHNet and three literature comparator methods: PIGNet, GAABind, and taqo [18-20]. The benchmarking analysis was designed to evaluate the ability of protein-ligand interaction scoring models to discriminate experimentally active ligands from property-matched decoys for a given target. Comparator methods were selected to represent distinct contemporary modelling paradigms, including GAABind, representing geometry-aware attention-based neural architectures; PIGNet, representing physics-informed graph neural-network approaches; and taqo, representing scoring based on machine-learned interatomic potentials. These methods were selected because they provide protein-ligand interaction scores that can be evaluated directly for active-decoy discrimination and have demonstrated competitive performance in their respective published studies. Because the comparator methods produce scores on different numerical scales a KDE-based analysis was used to characterize the separation of active and inactive score distributions: separate kernel-density estimates were constructed for the two classes to create a model-specific discrimination threshold for active and inactive classification and the accuracies obtained are shared in Table 3 below.

**Figure 3.**
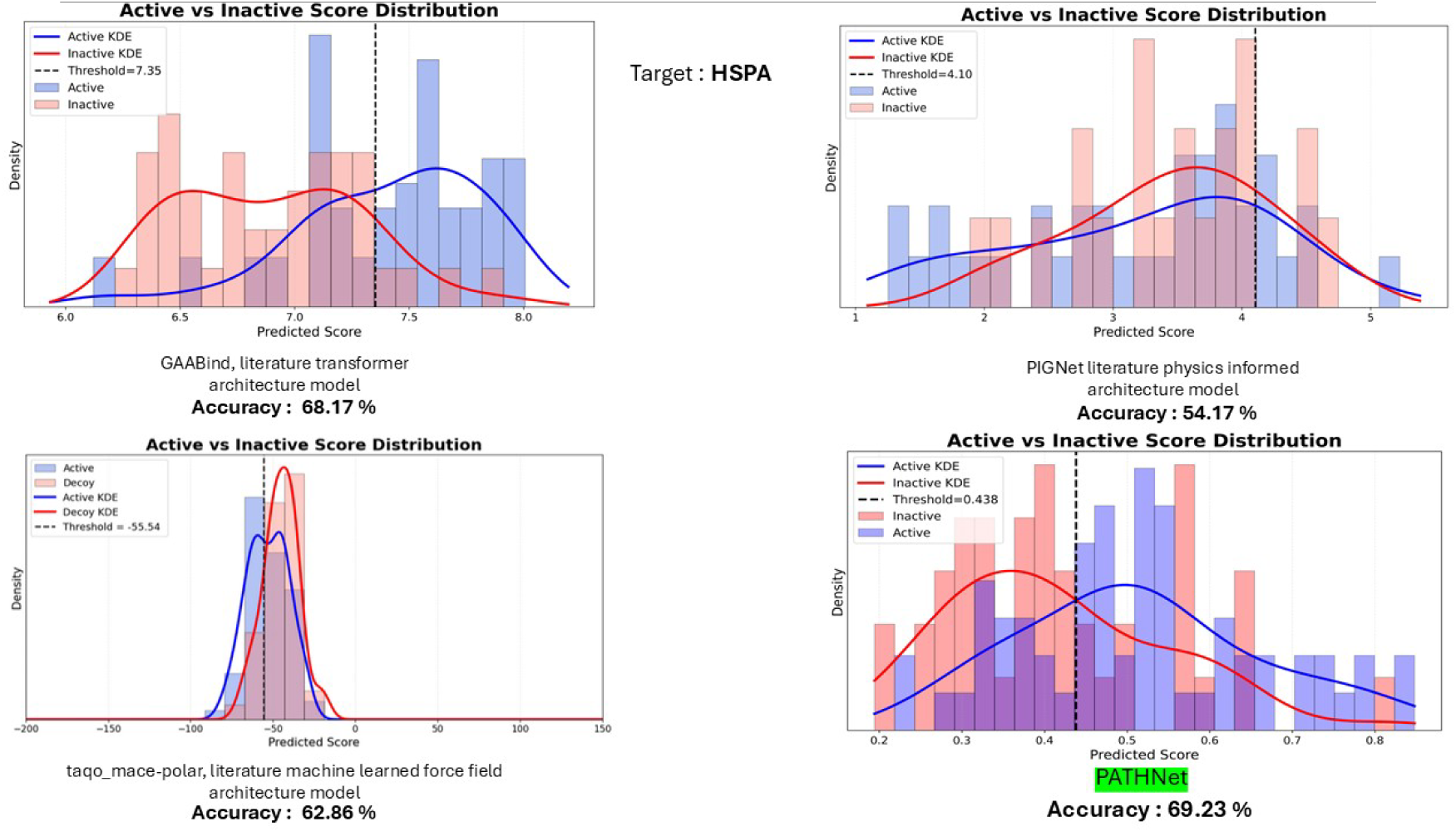
Supplied DUD-E validation panel for HSPA: active/inactive score distributions and reported accuracies for GAABind, PIGNet, taqo, and PATHNet

**Figure 4.**
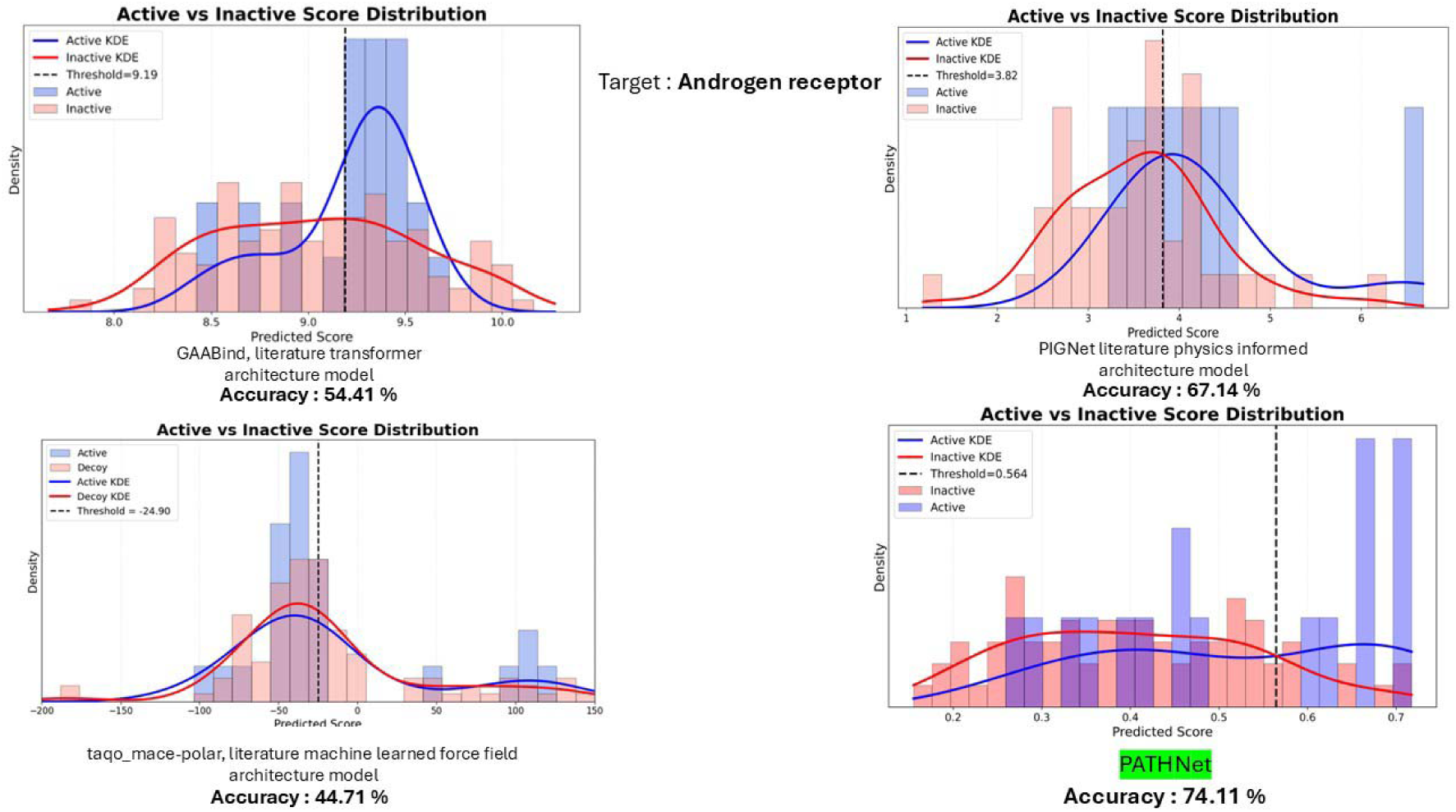
Supplied DUD-E validation panel for Androgen receptor: active/inactive score distributions and reported accuracies for GAABind, PIGNet, taqo, and PATHNet.

**Figure 5.**
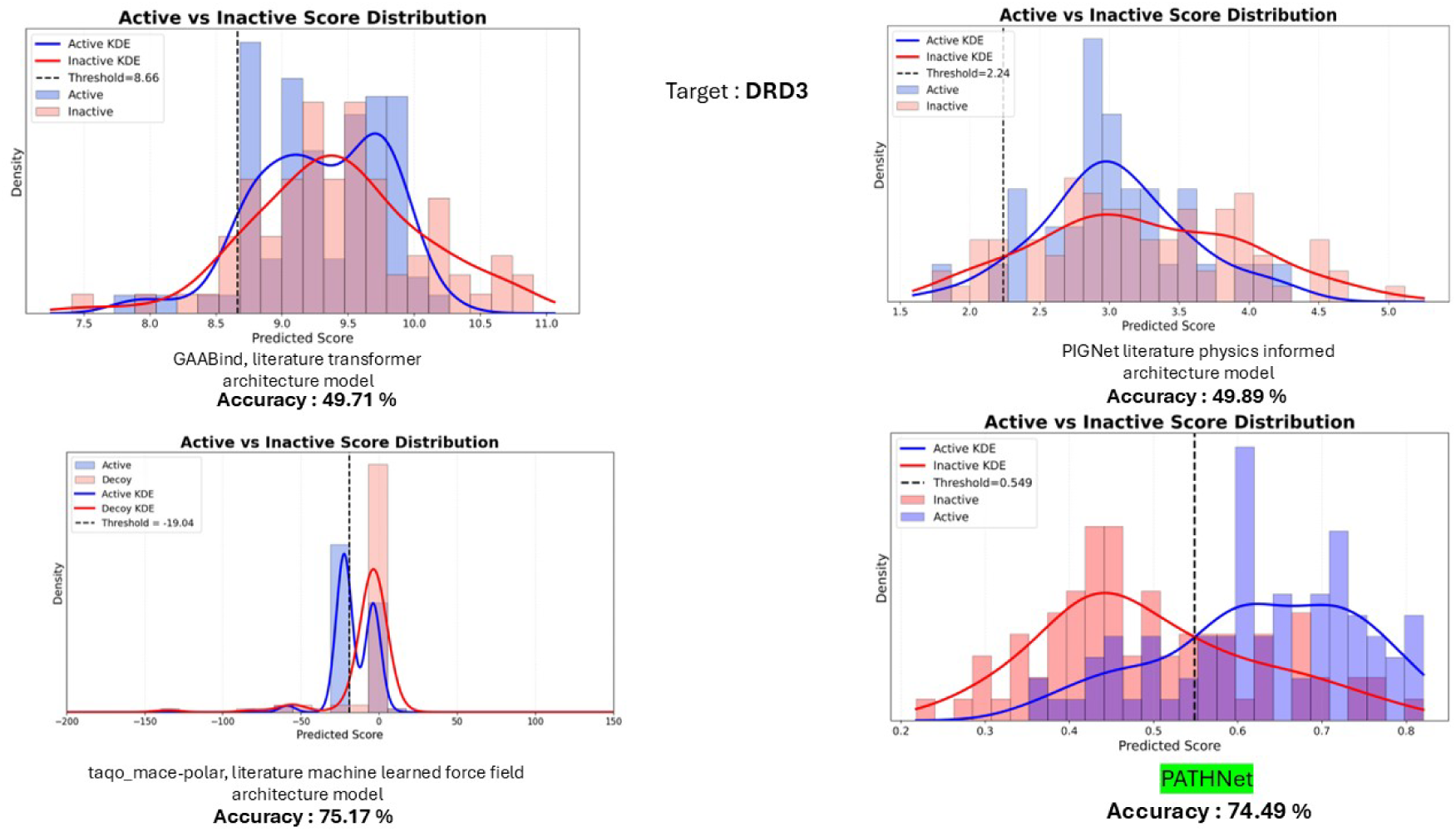
Supplied DUD-E validation panel for DRD3: active/inactive score distributions and reported accuracies for GAABind, PIGNet, taqo, and PATHNet

**Figure 6.**
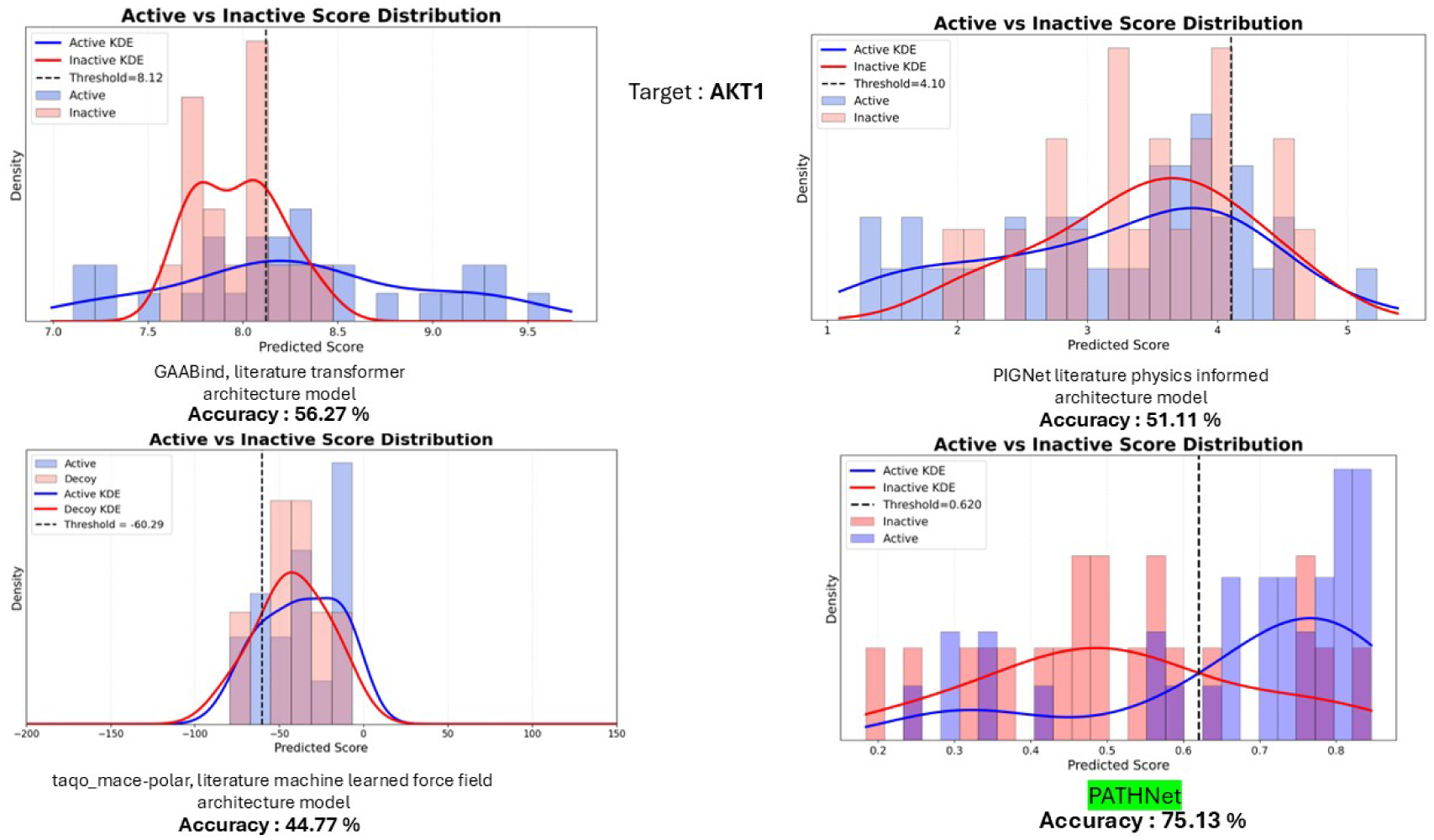
Supplied DUD-E validation panel for AKT1 (Serine/threonine-protein kinase AKT): active/inactive score distributions and reported accuracies for GAABind, PIGNet, taqo, and PATHNet

**Figure 7.**
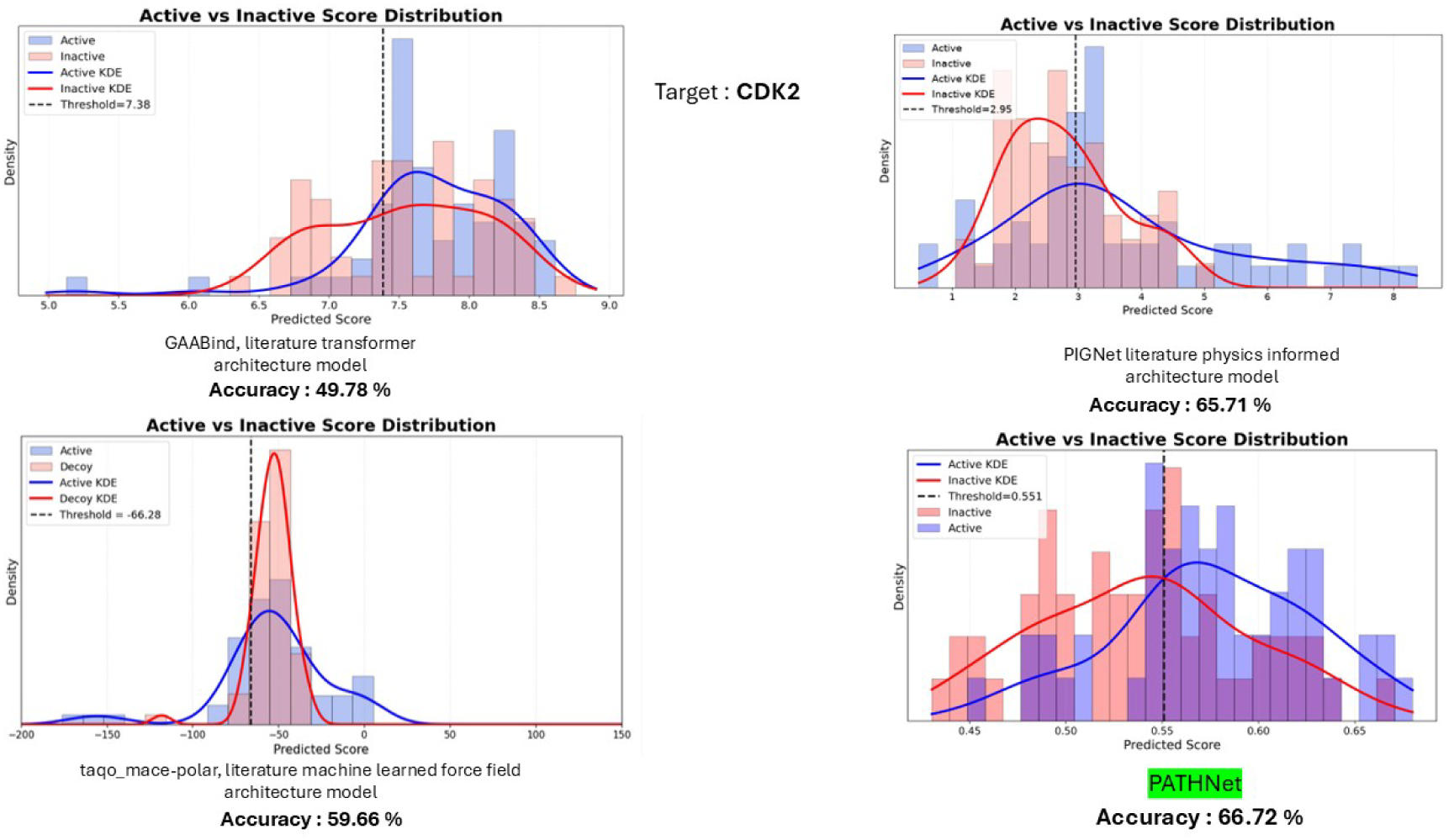
Supplied DUD-E validation panel for CDK2: active/inactive score distributions and reported accuracies for GAABind, PIGNet, taqo, and PATHNet

**Figure 8.**
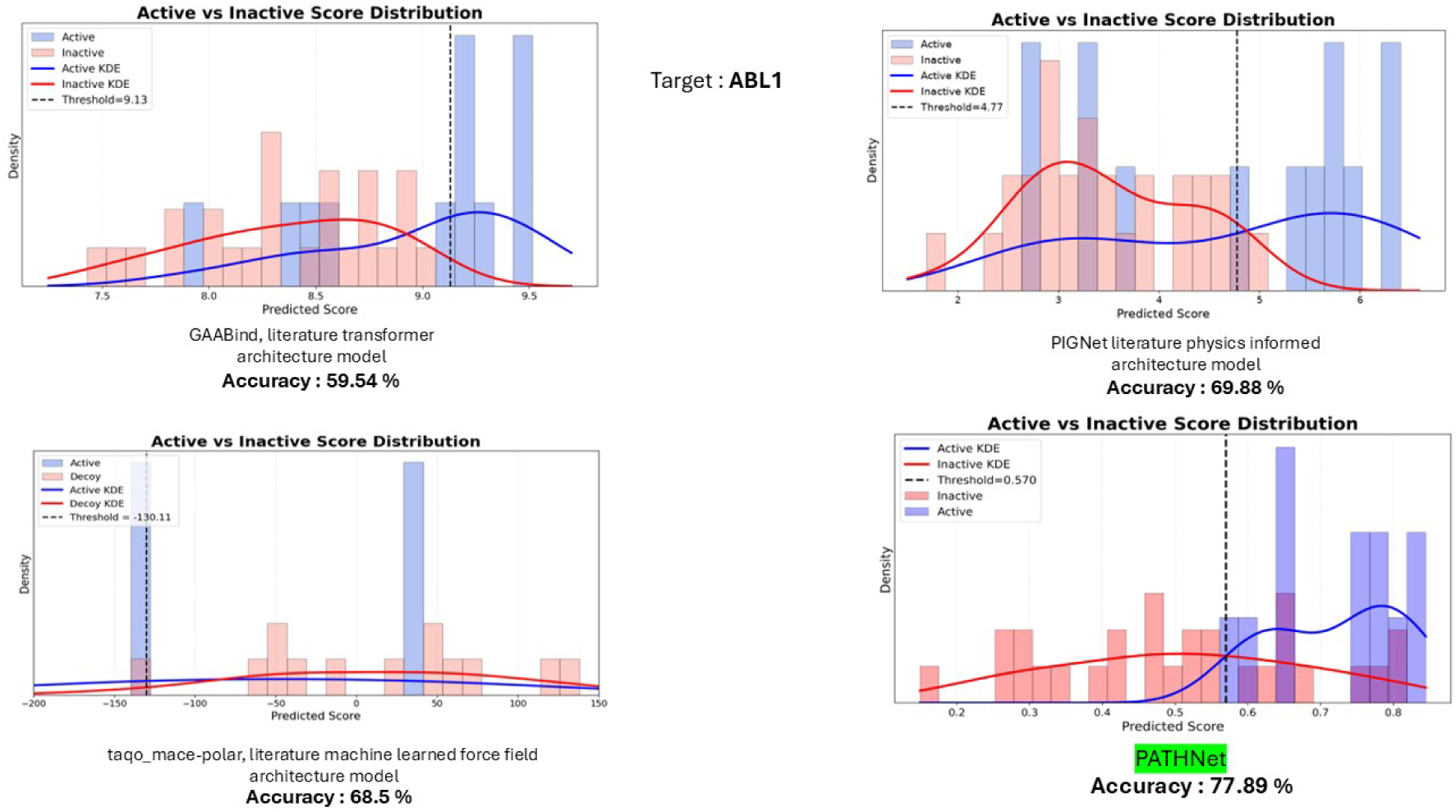
Supplied DUD-E validation panel for ABL1: active/inactive score distributions and reported accuracies for GAABind, PIGNet, taqo, and PATHNet

**Figure 9.**
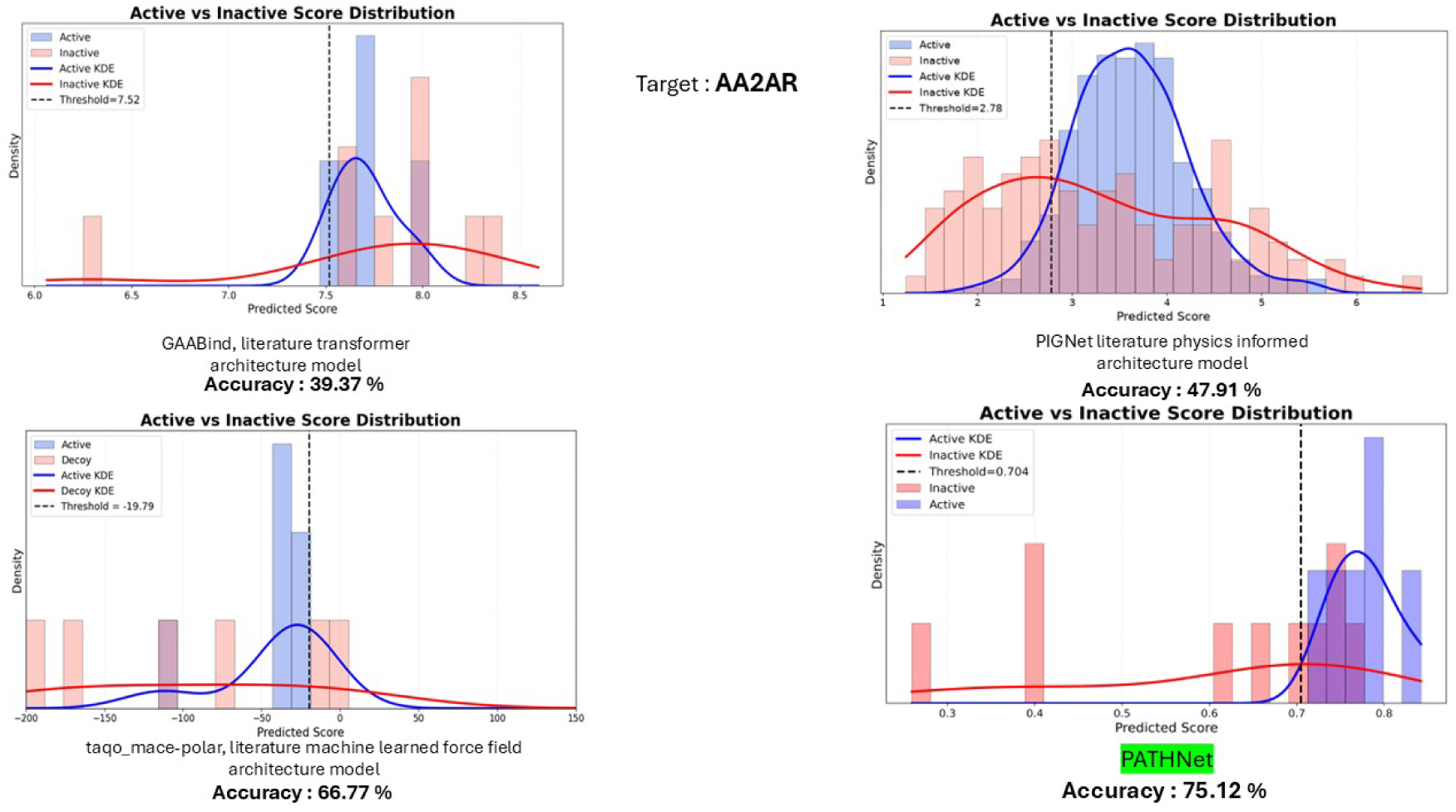
Supplied DUD-E validation panel for AA2AR (Adenosine A2A receptor): active/inactive score distributions and reported accuracies for GAABind, PIGNet, taqo, and PATHNet

**Table 3.** Reported classification accuracy (%) on the seven supplied DUD-E validation panels, including GAABind, PIGNet, taqo (MACE-POLAR), and PATHNet. The final column reports the PATHNet difference relative to the strongest comparator on each target.

| Target | GAABind | PIGNet | taqo<br>(MACE-POLAR) | PATHNet | $\Delta$ vs. best<br>baseline |
| --- | --- | --- | --- | --- | --- |
| HSPA | 68.17 | 54.17 | 62.86 | **69.23** | +1.06 |
| Androgen receptor | 54.41 | 67.14 | 44.71 | **74.11** | +6.97 |
| DRD3 | 49.71 | 49.89 | 75.17 | **74.49** | -0.68 |
| AKT1<br>(Serine/threonine-protein kinase AKT) | 56.27 | 51.11 | 44.77 | **75.13** | +18.86 |
| CDK2 | 49.78 | 65.71 | 59.66 | **66.72** | +1.01 |
| ABL1 | 59.54 | 69.88 | 68.50 | **77.89** | +8.01 |
| AA2AR<br>(Adenosine A2A receptor) | 39.37 | 47.91 | 66.77 | **75.12** | +8.35 |
| **Mean** | **53.89** | **57.97** | **60.35** | **73.24** | **+6.23** |

## Discussion

The multi-start optimization results indicate that the RG-inspired regulator changes not only the individual short-screen minima but also the quality of the solutions reached after extended optimization. Across the ten retained basins, the regulated runs consistently access lower-loss regions of parameter space, with short-screen minima extending down to 0.43 compared with 0.57 without regulation. More importantly, after all retained basins were subjected to extended optimization, the lowest converged loss decreased from 0.55 without regulation to 0.37 with regulation. This indicates that the regulator does not simply improve a single initialization but instead reshapes the accessible optimization landscape in a manner that promotes convergence toward lower-loss basins across the ensemble of candidate solutions. The result therefore supports the role of RG-inspired spectral regulation as a mechanism for stabilizing parameter evolution and improving access to more favourable regions of the non-convex loss landscape Across the seven panels, PATHNet achieved a highest mean reported accuracy of 73.24%, compared with 53.89% for GAABind, 57.97% for PIGNet, and 60.35% for taqo [18-20]. These targets also span distinct structural classes, including the molecular chaperone HSP90A, the kinases CDK2, AKT1, and ABL1, the class A GPCRs DRD3 and AA2AR, and the nuclear receptor AR. These proteins differ substantially in fold, pocket geometry, solvent exposure, conformational flexibility, and the balance of electrostatic, hydrophobic, and steric interactions governing ligand recognition. The higher mean performance and leading accuracy across six structurally heterogeneous targets therefore suggest that PATHNet is not restricted to a single binding-site topology and can generalize across diverse protein-ligand interaction environments.

Overall, these results support the broader interpretation of PATHNet as a physics-informed variational learning framework rather than a purely empirical scoring model. In the physical formulation, equilibrium and least-action solutions are obtained by searching over admissible paths until a stationary free-energy or action condition is reached; in PATHNet, the analogous search occurs in model-parameter space, where different parameter trajectories define competing learned functionals F_θ_(*X*) and convergence is identified by ∇*_θ_L*(θ*) = 0. The multi-start results show that RG-inspired regulation can improve access to lower-loss stationary basins, while the cross-target validation indicates that the resulting learned functional remains useful across structurally distinct binding environments. Combining physically meaningful descriptors, an explicit nonlinear functional mapper, a bounded neural correction, and regulated variational optimization therefore offers a practical route for learning transferable approximations to protein-ligand binding relationships, while preserving a clear correspondence between the optimization problem in the neural network and the variational structure of the underlying physics.

## 4. Conclusion, Limitations and Future Scope

PATHNet presents a physics-informed framework for protein-ligand binding prediction in which physically meaningful descriptors are combined with a flexible learned functional representation and a regulated optimization strategy. Instead of relying on a purely empirical mapping from generic molecular descriptors to activity, PATHNet learns the nonlinear relationship between non-bonded interactions, solvation, conformational strain, and the binding endpoint through an interpretable NM-PWL mapper supplemented by a bounded neural-network correction. The multi-start optimization results further indicate that RG-inspired regulation can guide the model toward lower-loss stationary solutions across multiple parameter basins, while validation across HSP90A, AR, DRD3, AKT1, CDK2, ABL1, and AA2AR demonstrates transfer across structurally and mechanistically diverse protein targets. PATHNet achieved a mean reported accuracy of 73.24%, compared with 53.89% for GAABind, 57.97% for PIGNet, and 60.35% for taqo. The broad cross-target performance is consistent with the physics-based grounding of PATHNet, in which transferable electrostatic, van der Waals, solvation, and conformational-strain quantities anchor the learned representation across different protein environments. These findings support the use of variationally motivated, physics-grounded neural architectures as a practical route toward more transferable protein-ligand scoring and pose-ranking models, while retaining a clear correspondence between stationarity in the underlying physical problem and convergence in neural-network parameter space.

The principal limitations of the present study arise from the restricted number of receptor families and ligand scaffolds represented in the current validation set, together with physical contributions that are not yet explicitly incorporated into the learned binding functional. In particular, effects arising from explicit water rearrangement, solvent reorganization, and the associated entropic contributions are not directly represented in the present feature set. Future work will therefore focus on extending validation across broader receptor families and chemotypes and on incorporating these additional physical terms, which should further improve the completeness, robustness, and transferability of the PATHNet framework.

## Conflict of Interest

The authors declare no competing financial interest.

## Data and Code Availability

The data and code required to reproduce the results reported in this manuscript are provided in the supplementary file and their description is as follows:

## Dataset: PDBbind

PDBbind complexes with reported ligand potency of 1 µM or better were selected as the starting dataset.

**Supplementary filename:** PATHNet_PDBBind_set

## Active-decoy dataset generation: AutoDock Vina

The crystallographic ligand was redocked into its cognate receptor using AutoDock Vina. Poses with heavy-atom RMSD <3 Å from the crystal structure were classified as native-like/active, while other retained low-energy poses were used as decoys.

**Supplementary filename:** PATHNet_active_decoy_generation

## PATHNet physics-based feature generation

Non bonded interaction, solvation, and ligand conformational-strain descriptors were calculated for each protein-ligand pose using the supplied feature-generation scripts.

**Supplementary filename:** PATHNet_feature_generation

## PATHNet modelling

The calculated physics descriptors were used to train PATHNet using the supplied architecture and optimization workflow, including feature weighting, the NM-PWL mapper, neural-network correction, and regulated optimization.

**Supplementary filename:** PATHNet_model/ RG_regulator

## External validation: DUD-E

External validation was carried out on HSP90A ( https://dude.docking.org/targets/hs90a), androgen receptor (AR) ( https://dude.docking.org/targets/andr), DRD3 ( https://dude.docking.org/targets/drd3), AKT1 ( https://dude.docking.org/targets/akt1), CDK2 ( https://dude.docking.org/targets/cdk2), ABL1 ( https://dude.docking.org/targets/abl1), and AA2AR ( https://dude.docking.org/targets/aa2ar) using the respective actives and decoys data deposited against each target in the DUD-E database.

Together with the methodology described in the manuscript, these resources provide the complete path for reproducing the reported results.

## Author Contributions

**Conceptualization and methodology:** Ben Geoffrey A. S., Abhishek Singh, Sowmya Kanchan, conceived the study, developed the research framework, and contributed to the overall architecture and design of the work.

**Model development and implementation:** Abhishek Singh, Ben Geoffrey A. S., and Sowmya Kanchan developed, implemented, and optimized the computational models used in this study.

**Data analysis and interpretation:** Ben Geoffrey A. S., Abhishek Singh, and Sowmya Kanchan analysed the results, interpreted the findings, and contributed to the scientific discussion of the outcomes.

**Writing:** Original Draft Preparation: Nagaraj M. Kulkarni and Ben Geoffrey A. S.

**Review & Editing:** All authors.

**Supervision and Project Administration:** Nagaraj M. Kulkarni.

## Notes

### Competing Interest Statement

The authors have declared no competing interest.

