## Supplementary material for "PATHNet: A Thermodynamic Path-Integral-Inspired Neural Network for Protein-Ligand Interaction Prediction": Step2_PATHNet_active_decoy_generation.py

```
#!/usr/bin/env python3

import argparse
import csv
import math
import os
from pathlib import Path
import shutil
import subprocess

import numpy as np
from rdkit import Chem
from rdkit.Chem import AllChem


def run(cmd, cwd=None):
    print(" ".join(map(str, cmd)))
    p = subprocess.run(
        list(map(str, cmd)),
        cwd=cwd,
        text=True,
        stdout=subprocess.PIPE,
        stderr=subprocess.STDOUT
    )
    if p.returncode != 0:
        raise RuntimeError(
            "Command failed:\n" + " ".join(map(str, cmd)) + "\n\n" + p.stdout
        )
    return p.stdout


def require_executables(names):
    missing = [x for x in names if shutil.which(x) is None]
    if missing:
        raise RuntimeError("Missing executables: " + ", ".join(missing))


def find_pdbbind_files(pdb_dir):
    pdb_id = pdb_dir.name

    protein_candidates = [
        pdb_dir / f"{pdb_id}_protein.pdb",
        pdb_dir / "protein.pdb",
    ]
    ligand_candidates = [
        pdb_dir / f"{pdb_id}_ligand.mol2",
        pdb_dir / f"{pdb_id}_ligand.sdf",
        pdb_dir / "ligand.mol2",
        pdb_dir / "ligand.sdf",
    ]

    protein = next((x for x in protein_candidates if x.exists()), None)
    ligand = next((x for x in ligand_candidates if x.exists()), None)

    return protein, ligand


def load_reference_ligand(path):
    suffix = path.suffix.lower()

    if suffix == ".mol2":
        mol = Chem.MolFromMol2File(str(path), removeHs=False)
    elif suffix == ".sdf":
        supplier = Chem.SDMolSupplier(str(path), removeHs=False)
        mol = next((m for m in supplier if m is not None), None)
    elif suffix == ".mol":
        mol = Chem.MolFromMolFile(str(path), removeHs=False)
    else:
        raise ValueError(f"Unsupported ligand format: {suffix}")

    if mol is None:
        raise RuntimeError(f"RDKit could not read {path}")

    return mol


def ligand_box(reference_mol, margin=12.0, minimum=22.0, maximum=32.0):
    mol = Chem.RemoveHs(reference_mol)
    conf = mol.GetConformer()

    xyz = np.array([
        list(conf.GetAtomPosition(i))
        for i in range(mol.GetNumAtoms())
    ], dtype=float)

    center = 0.5 * (xyz.min(axis=0) + xyz.max(axis=0))
    extent = xyz.max(axis=0) - xyz.min(axis=0)

    size = np.clip(extent + float(margin), float(minimum), float(maximum))

    return center, size


def prepare_receptor(protein_pdb, receptor_pdbqt, mgl_python, prepare_receptor):
    run([
        mgl_python,
        prepare_receptor,
        "-A", "bonds_hydrogens",
        "-U", "nphs_lps_waters_nonstdres_deleteAltB",
        "-r", str(protein_pdb),
        "-o", str(receptor_pdbqt),
    ])


def prepare_ligand(ligand_file, workdir, ligand_pdbqt,
                   mgl_python, prepare_ligand):
    ligand_pdb = workdir / "ligand_reference.pdb"

    run([
        "obabel",
        str(ligand_file),
        "-opdb",
        "-O", str(ligand_pdb),
    ])

    run([
        mgl_python,
        prepare_ligand,
        "-A", "bonds_hydrogens",
        "-U", "nphs_lps",
        "-l", str(ligand_pdb),
        "-o", str(ligand_pdbqt),
    ])


def run_vina(receptor_pdbqt, ligand_pdbqt, out_pdbqt, log_file,
             center, size, exhaustiveness, num_modes, energy_range, seed=None):
    cmd = [
        "vina",
        "--receptor", str(receptor_pdbqt),
        "--ligand", str(ligand_pdbqt),
        "--center_x", f"{center[0]:.4f}",
        "--center_y", f"{center[1]:.4f}",
        "--center_z", f"{center[2]:.4f}",
        "--size_x", f"{size[0]:.4f}",
        "--size_y", f"{size[1]:.4f}",
        "--size_z", f"{size[2]:.4f}",
        "--exhaustiveness", str(exhaustiveness),
        "--num_modes", str(num_modes),
        "--energy_range", str(energy_range),
        "--out", str(out_pdbqt),
        "--log", str(log_file),
    ]

    if seed is not None:
        cmd += ["--seed", str(seed)]

    run(cmd)


def split_vina_models(vina_out, pose_dir):
    pose_dir.mkdir(parents=True, exist_ok=True)

    models = []
    current = []
    model_id = None
    affinity = None

    with open(vina_out) as fh:
        for line in fh:
            if line.startswith("MODEL"):
                current = [line]
                parts = line.split()
                model_id = int(parts[1]) if len(parts) > 1 else len(models) + 1
                affinity = None

            elif line.startswith("REMARK VINA RESULT:"):
                current.append(line)
                parts = line.split()
                affinity = float(parts[3])

            elif line.startswith("ENDMDL"):
                current.append(line)

                pose_file = pose_dir / f"pose_{model_id:03d}.pdbqt"
                pose_file.write_text("".join(current))

                models.append({
                    "model": model_id,
                    "affinity": affinity,
                    "pdbqt": pose_file,
                })

                current = []
                model_id = None
                affinity = None

            elif current:
                current.append(line)

    if not models:
        raise RuntimeError(f"No Vina models found in {vina_out}")

    return models


def pose_pdbqt_to_mol(pose_pdbqt, reference_mol, workdir):
    pose_pdb = workdir / (pose_pdbqt.stem + ".pdb")

    run([
        "obabel",
        str(pose_pdbqt),
        "-opdb",
        "-O", str(pose_pdb),
    ])

    pose = Chem.MolFromPDBFile(
        str(pose_pdb),
        removeHs=False,
        sanitize=False
    )

    if pose is None:
        raise RuntimeError(f"RDKit could not read converted pose {pose_pdb}")

    ref_noh = Chem.RemoveHs(reference_mol)
    pose_noh = Chem.RemoveHs(pose, sanitize=False)

    try:
        pose_typed = AllChem.AssignBondOrdersFromTemplate(ref_noh, pose_noh)
    except Exception:
        pose_typed = pose_noh

    Chem.SanitizeMol(pose_typed)

    return pose_typed


def direct_symmetry_aware_rmsd(reference_mol, pose_mol, max_matches=100000):
    ref = Chem.RemoveHs(reference_mol)
    pose = Chem.RemoveHs(pose_mol)

    if ref.GetNumAtoms() != pose.GetNumAtoms():
        raise RuntimeError(
            f"Atom-count mismatch: reference={ref.GetNumAtoms()}, "
            f"pose={pose.GetNumAtoms()}"
        )

    ref_conf = ref.GetConformer()
    pose_conf = pose.GetConformer()

    ref_xyz = np.array([
        list(ref_conf.GetAtomPosition(i))
        for i in range(ref.GetNumAtoms())
    ], dtype=float)

    matches = pose.GetSubstructMatches(
        ref,
        uniquify=False,
        useChirality=False,
        maxMatches=max_matches
    )

    if not matches:
        match = pose.GetSubstructMatch(ref, useChirality=False)
        matches = [match] if match else []

    if not matches:
        raise RuntimeError("Could not map docked pose atoms to reference ligand.")

    best = math.inf

    for match in matches:
        pose_xyz = np.array([
            list(pose_conf.GetAtomPosition(match[i]))
            for i in range(ref.GetNumAtoms())
        ], dtype=float)

        rmsd = float(np.sqrt(np.mean(np.sum(
            (ref_xyz - pose_xyz) ** 2,
            axis=1
        ))))

        if rmsd < best:
            best = rmsd

    return best


def molecule_to_sdf(mol, out_sdf):
    writer = Chem.SDWriter(str(out_sdf))
    writer.write(mol)
    writer.close()


def low_energy_pose_records(models, reference_mol, pose_dir, energy_window):
    best_affinity = min(
        m["affinity"] for m in models
        if m["affinity"] is not None
    )
    cutoff = best_affinity + float(energy_window)

    records = []

    for m in models:
        if m["affinity"] is None:
            continue
        if m["affinity"] > cutoff:
            continue

        mol = pose_pdbqt_to_mol(
            m["pdbqt"],
            reference_mol,
            pose_dir
        )

        rmsd_to_ref = direct_symmetry_aware_rmsd(
            reference_mol,
            mol
        )

        records.append({
            **m,
            "mol": mol,
            "rmsd_to_crystal": rmsd_to_ref,
        })

    records.sort(key=lambda x: x["affinity"])
    return records, best_affinity, cutoff


def cluster_low_energy_poses(records, cluster_rmsd):
    representatives = []

    for record in records:
        belongs_to_existing = False

        for rep in representatives:
            rmsd = direct_symmetry_aware_rmsd(
                rep["mol"],
                record["mol"]
            )

            if rmsd < float(cluster_rmsd):
                belongs_to_existing = True
                break

        if not belongs_to_existing:
            representatives.append(record)

    return representatives


def save_classified_pose(record, out_dir, pdb_id, label, cluster_index):
    out_dir.mkdir(parents=True, exist_ok=True)

    out_sdf = out_dir / (
        f"{pdb_id}_{label}_cluster{cluster_index:02d}_"
        f"vina{record['affinity']:.2f}_"
        f"rmsd{record['rmsd_to_crystal']:.2f}.sdf"
    )

    molecule_to_sdf(record["mol"], out_sdf)
    return out_sdf


def process_target(pdb_dir, output_root, args, writer):
    pdb_id = pdb_dir.name
    protein, ligand = find_pdbbind_files(pdb_dir)

    if protein is None or ligand is None:
        writer.writerow({
            "pdb_id": pdb_id,
            "status": "missing_input",
        })
        return

    target_out = output_root / pdb_id
    active_dir = target_out / "actives"
    decoy_dir = target_out / "decoys"
    workdir = target_out / "_vina_work"

    active_dir.mkdir(parents=True, exist_ok=True)
    decoy_dir.mkdir(parents=True, exist_ok=True)
    workdir.mkdir(parents=True, exist_ok=True)

    shutil.copy2(protein, target_out / "protein.pdb")
    shutil.copy2(ligand, target_out / ligand.name)

    reference_mol = load_reference_ligand(ligand)
    center, size = ligand_box(
        reference_mol,
        margin=args.box_margin,
        minimum=args.box_min,
        maximum=args.box_max
    )

    receptor_pdbqt = workdir / "receptor.pdbqt"
    ligand_pdbqt = workdir / "ligand.pdbqt"
    vina_out = workdir / "vina_out.pdbqt"
    vina_log = workdir / "vina.log"
    pose_dir = workdir / "poses"

    prepare_receptor(
        protein,
        receptor_pdbqt,
        args.mgl_python,
        args.prepare_receptor
    )

    prepare_ligand(
        ligand,
        workdir,
        ligand_pdbqt,
        args.mgl_python,
        args.prepare_ligand
    )

    run_vina(
        receptor_pdbqt,
        ligand_pdbqt,
        vina_out,
        vina_log,
        center,
        size,
        args.exhaustiveness,
        args.num_modes,
        args.energy_range,
        args.seed
    )

    models = split_vina_models(vina_out, pose_dir)

    low_energy, best_affinity, energy_cutoff = low_energy_pose_records(
        models,
        reference_mol,
        pose_dir,
        args.low_energy_window
    )

    representatives = cluster_low_energy_poses(
        low_energy,
        args.cluster_rmsd
    )

    n_active = 0
    n_decoy = 0

    for cluster_index, record in enumerate(representatives, start=1):
        if record["rmsd_to_crystal"] < args.active_rmsd:
            label = "active"
            out_dir = active_dir
            n_active += 1
        else:
            label = "decoy"
            out_dir = decoy_dir
            n_decoy += 1

        saved = save_classified_pose(
            record,
            out_dir,
            pdb_id,
            label,
            cluster_index
        )

        writer.writerow({
            "pdb_id": pdb_id,
            "status": "ok",
            "cluster": cluster_index,
            "vina_model": record["model"],
            "vina_affinity_kcal_mol": record["affinity"],
            "best_vina_affinity_kcal_mol": best_affinity,
            "low_energy_cutoff_kcal_mol": energy_cutoff,
            "rmsd_to_crystal_A": record["rmsd_to_crystal"],
            "class": label,
            "saved_pose": str(saved),
        })

    if n_active == 0:
        writer.writerow({
            "pdb_id": pdb_id,
            "status": "redocking_failed_no_pose_below_rmsd_threshold",
            "best_vina_affinity_kcal_mol": best_affinity,
            "low_energy_cutoff_kcal_mol": energy_cutoff,
        })

    print(
        f"{pdb_id}: low-energy poses={len(low_energy)}, "
        f"clusters={len(representatives)}, "
        f"actives={n_active}, decoys={n_decoy}"
    )


def main():
    ap = argparse.ArgumentParser(
        description=(
            "PDBbind AutoDock Vina redocking pipeline: "
            "cluster low-energy poses, retain <3 A heavy-atom RMSD "
            "cluster representatives as active/native-like poses, "
            "and classify other low-energy cluster representatives as decoys."
        )
    )

    ap.add_argument("--pdbbind-root", required=True)
    ap.add_argument("--output-root", default="pdbbind_vina_redocking")
    ap.add_argument("--summary", default="vina_redocking_summary.csv")

    ap.add_argument(
        "--mgl-python",
        default=os.path.expanduser("~/MGLTools-1.5.6/bin/pythonsh")
    )
    ap.add_argument(
        "--prepare-receptor",
        default=os.path.expanduser(
            "~/MGLTools-1.5.6/MGLToolsPckgs/"
            "AutoDockTools/Utilities24/prepare_receptor4.py"
        )
    )
    ap.add_argument(
        "--prepare-ligand",
        default=os.path.expanduser(
            "~/MGLTools-1.5.6/MGLToolsPckgs/"
            "AutoDockTools/Utilities24/prepare_ligand4.py"
        )
    )

    ap.add_argument("--active-rmsd", type=float, default=3.0)
    ap.add_argument("--cluster-rmsd", type=float, default=2.0)
    ap.add_argument("--low-energy-window", type=float, default=3.0)

    ap.add_argument("--exhaustiveness", type=int, default=32)
    ap.add_argument("--num-modes", type=int, default=20)
    ap.add_argument("--energy-range", type=float, default=5.0)
    ap.add_argument("--seed", type=int, default=2026)

    ap.add_argument("--box-margin", type=float, default=12.0)
    ap.add_argument("--box-min", type=float, default=22.0)
    ap.add_argument("--box-max", type=float, default=32.0)

    ap.add_argument(
        "--pdb-ids",
        nargs="*",
        default=None,
        help="Optional subset of PDB IDs; otherwise process all directories."
    )

    args = ap.parse_args()

    require_executables(["vina", "obabel"])

    if not Path(args.mgl_python).exists():
        raise RuntimeError(f"MGLTools pythonsh not found: {args.mgl_python}")
    if not Path(args.prepare_receptor).exists():
        raise RuntimeError(
            f"prepare_receptor4.py not found: {args.prepare_receptor}"
        )
    if not Path(args.prepare_ligand).exists():
        raise RuntimeError(
            f"prepare_ligand4.py not found: {args.prepare_ligand}"
        )

    pdbbind_root = Path(args.pdbbind_root).resolve()
    output_root = Path(args.output_root).resolve()
    output_root.mkdir(parents=True, exist_ok=True)

    if args.pdb_ids:
        pdb_dirs = [
            pdbbind_root / pdb_id
            for pdb_id in args.pdb_ids
        ]
    else:
        pdb_dirs = sorted(
            x for x in pdbbind_root.iterdir()
            if x.is_dir()
        )

    fields = [
        "pdb_id",
        "status",
        "cluster",
        "vina_model",
        "vina_affinity_kcal_mol",
        "best_vina_affinity_kcal_mol",
        "low_energy_cutoff_kcal_mol",
        "rmsd_to_crystal_A",
        "class",
        "saved_pose",
    ]

    with open(args.summary, "w", newline="") as fh:
        writer = csv.DictWriter(fh, fieldnames=fields)
        writer.writeheader()

        for pdb_dir in pdb_dirs:
            try:
                process_target(
                    pdb_dir,
                    output_root,
                    args,
                    writer
                )
                fh.flush()

            except Exception as exc:
                print(f"FAILED {pdb_dir.name}: {exc}")

                writer.writerow({
                    "pdb_id": pdb_dir.name,
                    "status": "failed",
                    "saved_pose": str(exc),
                })
                fh.flush()


if __name__ == "__main__":
    main()
```
