## Supplementary material for "PATHNet: A Thermodynamic Path-Integral-Inspired Neural Network for Protein-Ligand Interaction Prediction": Step4_PATHNet_model.py

```
import tensorflow as tf
from tensorflow.keras import layers, Model


# ============================================================
# Piecewise-linear functional mapper
# ============================================================
class PiecewiseLinearMapper(layers.Layer):
    """
    Trainable piecewise-linear function:
        z in [0,1] --> f_map(z) in [0,1]

    Signed increments allow the learned functional form
    to be increasing, decreasing or non-monotonic.
    """

    def __init__(self, K=11, **kwargs):
        super().__init__(**kwargs)
        self.K = K

    def build(self, input_shape):

        self.knot_x = tf.linspace(0.0, 1.0, self.K)

        # Trainable values defining the learned function
        self.raw_y0 = self.add_weight(
            name="raw_y0",
            shape=(),
            initializer="zeros",
            trainable=True
        )

        self.raw_increments = self.add_weight(
            name="raw_increments",
            shape=(self.K - 1,),
            initializer=tf.keras.initializers.Constant(
                1.0 / (self.K - 1)
            ),
            trainable=True
        )

    def call(self, z):

        z = tf.clip_by_value(z, 0.0, 1.0)

        # Construct function values at knots
        y = self.raw_y0 + tf.concat(
            [
                tf.zeros((1,), dtype=z.dtype),
                tf.cumsum(self.raw_increments)
            ],
            axis=0
        )

        # Normalize learned mapper to [0,1]
        ymin = tf.reduce_min(y)
        ymax = tf.reduce_max(y)

        y = (y - ymin) / (ymax - ymin + 1e-8)

        # Linear interpolation between knots
        z_flat = tf.reshape(z, [-1])

        idx = tf.searchsorted(
            self.knot_x,
            z_flat,
            side="right"
        )

        idx = tf.clip_by_value(
            idx,
            1,
            self.K - 1
        )

        i0 = idx - 1
        i1 = idx

        x0 = tf.gather(self.knot_x, i0)
        x1 = tf.gather(self.knot_x, i1)

        y0 = tf.gather(y, i0)
        y1 = tf.gather(y, i1)

        t = (z_flat - x0) / (x1 - x0 + 1e-8)

        output = y0 + t * (y1 - y0)

        return tf.reshape(output, tf.shape(z))


# ============================================================
# PATHnet architecture snippet
# ============================================================
class PATHNetCore(Model):

    def __init__(
        self,
        mapper_knots=11,
        alpha_max=0.20
    ):

        super().__init__()

        # ----------------------------------------------------
        # Trainable feature importance
        #
        # Three physics-derived features:
        #   0 = non-bonded interaction
        #   1 = solvation
        #   2 = conformational strain
        # ----------------------------------------------------
        self.feature_logits = self.add_weight(
            name="physics_feature_logits",
            shape=(3,),
            initializer="zeros",
            trainable=True
        )

        # ----------------------------------------------------
        # Piecewise functional mapper
        # ----------------------------------------------------
        self.mapper = PiecewiseLinearMapper(
            K=mapper_knots,
            name="binding_function_mapper"
        )

        # ----------------------------------------------------
        # Neural-network correction
        # ----------------------------------------------------
        self.nn_hidden1 = layers.Dense(
            16,
            activation="tanh"
        )

        self.nn_hidden2 = layers.Dense(
            8,
            activation="tanh"
        )

        # sigmoid bounds neural output to [0,1]
        self.nn_output = layers.Dense(
            1,
            activation="sigmoid"
        )

        # ----------------------------------------------------
        # Learnable bounded mapper / NN mixing
        # ----------------------------------------------------
        self.alpha_raw = self.add_weight(
            name="alpha_raw",
            shape=(),
            initializer=tf.keras.initializers.Constant(-3.0),
            trainable=True
        )

        self.alpha_max = alpha_max


    def call(self, x, return_components=False):
        """
        x[:,0] = normalized non-bonded interaction descriptor
        x[:,1] = normalized solvation descriptor
        x[:,2] = normalized ligand-strain descriptor
        """

        x = tf.cast(x, tf.float32)
        x = tf.clip_by_value(x, 0.0, 1.0)

        # ====================================================
        # 1. LEARN FEATURE IMPORTANCE THROUGH SOFTMAX
        # ====================================================

        weights = tf.nn.softmax(
            self.feature_logits
        )

        # w_i > 0 and sum_i w_i = 1
        #
        # z = w_nonbond * E_nonbond
        #   + w_solv    * E_solv
        #   + w_strain  * E_strain

        z = tf.reduce_sum(
            x * weights[None, :],
            axis=-1,
            keepdims=True
        )

        z = tf.clip_by_value(
            z,
            0.0,
            1.0
        )

        # ====================================================
        # 2. LEARN UNKNOWN FUNCTIONAL FORM
        # ====================================================

        f_map = self.mapper(z)

        # ====================================================
        # 3. BOUNDED NEURAL-NETWORK CORRECTION
        # ====================================================

        h = self.nn_hidden1(x)
        h = self.nn_hidden2(h)

        f_nn = self.nn_output(h)

        # ====================================================
        # 4. LEARNED BOUNDED CORRECTION TO MAPPER
        # ====================================================

        alpha = (
            self.alpha_max *
            tf.sigmoid(self.alpha_raw)
        )

        f_corr = (
            (1.0 - alpha) * f_map
            +
            alpha * f_nn
        )

        f_corr = tf.clip_by_value(
            f_corr,
            0.0,
            1.0
        )

        if return_components:

            return {
                "prediction": f_corr,
                "feature_weights": weights,
                "mixed_physics_coordinate": z,
                "mapper_output": f_map,
                "nn_correction": f_nn,
                "alpha": alpha
            }

        return f_corr
```
