## Supplementary material for "PATHNet: A Thermodynamic Path-Integral-Inspired Neural Network for Protein-Ligand Interaction Prediction": Step4_RG_regulator.py

```
from __future__ import annotations

import math
from typing import Dict, Optional

import torch
import torch.nn as nn


_SUPPORTED_MODULES = (nn.Linear, nn.Embedding)


def _is_supported_matrix(module: nn.Module) -> bool:

    if not isinstance(module, _SUPPORTED_MODULES):
        return False

    weight = getattr(module, "weight", None)
    return weight is not None and weight.ndim == 2 and min(weight.shape) >= 2


def _finite_scalar(value: Optional[float]) -> bool:

    if value is None:
        return False

    try:
        return math.isfinite(float(value))
    except (TypeError, ValueError):
        return False


def _copy_matrix_(module: nn.Module, matrix: torch.Tensor) -> None:

    if hasattr(module, "parametrizations") and "weight" in module.parametrizations:
        raise TypeError(
            "regulator_dmpk_mask expects plain nn.Linear/nn.Embedding weights; "
            "remove weight parametrization or add a parametrization-specific "
            "write-back rule."
        )

    module.weight.copy_(matrix)


@torch.no_grad()
def dynamic_band_spectral_regulate(
    module: nn.Module,
    layer_name: str,
    *,


    max_ratio: float = 2.0,
    min_ratio: float = 1.0,
    ratio_tolerance: float = 0.1,
    eta: float = 1.0,


    gamma: float = 1.0e-6,


    grad_max_ratio: Optional[float] = 2.0,
    grad_ratio_tolerance: float = 0.1,


    current_val_loss: Optional[float] = None,
    mask_after_val_loss: Optional[float] = None,
    mask_threshold: float = 0.1,


    eps: float = 1.0e-8,
    verbose: bool = False,
) -> Dict[str, object]:


    status: Dict[str, object] = {
        "layer": layer_name,
        "supported": False,
        "regulated": False,
        "mask_triggered": False,
        "mask_active": False,
        "masked_count": 0,
        "svd_failed": False,
    }

    if not _is_supported_matrix(module):
        return status

    if max_ratio < min_ratio:
        raise ValueError("Require min_ratio <= max_ratio.")
    if not (0.0 <= eta <= 1.0):
        raise ValueError("eta must be in [0, 1].")
    if not (0.0 < gamma <= 1.0):
        raise ValueError("gamma must be in (0, 1].")
    if not (0.0 <= mask_threshold <= 1.0):
        raise ValueError("mask_threshold must be in [0, 1].")
    if grad_max_ratio is not None and grad_max_ratio < 0.0:
        raise ValueError("grad_max_ratio must be non-negative or None.")

    status["supported"] = True
    weight_old = module.weight.detach().clone()


    try:
        U, singular_values, Vh = torch.linalg.svd(
            weight_old,
            full_matrices=False,
        )
    except Exception as exc:
        status["svd_failed"] = True
        if verbose:
            print(f"[Regulator] SVD failed on {layer_name}: {exc}")
        return status


    s_sorted, sort_idx = torch.sort(singular_values, descending=True)
    s_before = s_sorted.clone()
    s_target = s_sorted.clone()


    mask_active_before = bool(getattr(module, "_spectral_mask_active", False))
    trigger_condition = (
        not mask_active_before
        and _finite_scalar(current_val_loss)
        and _finite_scalar(mask_after_val_loss)
        and float(current_val_loss) <= float(mask_after_val_loss)
    )

    if trigger_condition:
        smax_mask = torch.clamp(s_target[0], min=eps)
        normalized = s_target / smax_mask
        keep_mask = normalized >= mask_threshold
        keep_mask[0] = True
        keep_count = max(1, int(keep_mask.sum().item()))

        module._spectral_mask_active = True
        module._spectral_keep_count = keep_count
        module._spectral_mask_threshold = float(mask_threshold)
        module._spectral_mask_val_loss = float(current_val_loss)
        module._spectral_prev_sorted = s_before.clone()

        status["mask_triggered"] = True

    mask_active = bool(getattr(module, "_spectral_mask_active", False))
    status["mask_active"] = mask_active

    if mask_active:
        keep_count = int(getattr(module, "_spectral_keep_count", len(s_target)))
        keep_count = max(1, min(keep_count, len(s_target)))

        if keep_count < len(s_target):
            s_target[keep_count:] = 0.0

        status["masked_count"] = len(s_target) - keep_count
    else:
        keep_count = len(s_target)


    active = s_target[:keep_count]

    log_ratio_before: Optional[float] = None
    log_ratio_after: Optional[float] = None
    frozen_ratio: Optional[float] = None
    correction_message = "spectrum healthy"


    if len(active) >= 2:
        smax = torch.clamp(active.max(), min=eps)
        smin = torch.clamp(active.min(), min=eps)

        log_ratio = torch.log(smax) - torch.log(smin)
        log_ratio_before = float(log_ratio.item())


        if mask_active and hasattr(module, "_spectral_ratio_target"):
            frozen_target = float(module._spectral_ratio_target)
            lower = max(1.0 + eps, frozen_target - ratio_tolerance)
            upper = max(lower, frozen_target + ratio_tolerance)
            raw_ratio = smax / smin

            if raw_ratio > upper:
                idx_min = torch.argmin(active)
                active[idx_min] = smax / upper
                correction_message = "post-mask frozen ratio corrected"
            elif raw_ratio < lower:
                idx_min = torch.argmin(active)
                active[idx_min] = smax / lower
                correction_message = "post-mask frozen ratio corrected"
        else:


            if log_ratio > (max_ratio + ratio_tolerance):
                target_smin = torch.exp(torch.log(smax) - max_ratio)
                idx_min = torch.argmin(active)
                active[idx_min] = (
                    (1.0 - eta) * active[idx_min]
                    + eta * target_smin
                )
                correction_message = "sharp spectrum corrected"


            elif log_ratio < (min_ratio - ratio_tolerance):
                target_smin = torch.exp(torch.log(smax) - min_ratio)
                idx_min = torch.argmin(active)
                active[idx_min] = (
                    (1.0 - eta) * active[idx_min]
                    + eta * target_smin
                )
                correction_message = "flat spectrum corrected"

        s_target[:keep_count] = active


        if status["mask_triggered"]:
            active_smax_after = torch.clamp(active.max(), min=eps)
            active_smin_after = torch.clamp(active.min(), min=eps)
            module._spectral_ratio_target = float(
                (active_smax_after / active_smin_after).item()
            )

        if hasattr(module, "_spectral_ratio_target"):
            frozen_ratio = float(module._spectral_ratio_target)

        smax_after = torch.clamp(active.max(), min=eps)
        smin_after = torch.clamp(active.min(), min=eps)
        log_ratio_after = float(
            (torch.log(smax_after) - torch.log(smin_after)).item()
        )


    s_target_unsorted = torch.empty_like(s_target)
    s_target_unsorted[sort_idx] = s_target


    if mask_active:
        previous_sorted = getattr(module, "_spectral_prev_sorted", s_before)

        if previous_sorted.shape != s_target.shape:
            previous_sorted = s_before

        s_applied_sorted = (
            previous_sorted
            + gamma * (s_target - previous_sorted)
        )
        module._spectral_prev_sorted = s_applied_sorted.detach().clone()

        s_applied = torch.empty_like(s_applied_sorted)
        s_applied[sort_idx] = s_applied_sorted
    else:
        s_applied = s_target_unsorted


    weight_candidate = (U * s_applied.unsqueeze(0)) @ Vh


    delta = weight_candidate - weight_old


    if not mask_active and grad_max_ratio is not None:
        abs_delta = delta.abs()
        valid_mask = abs_delta > eps

        if int(valid_mask.sum().item()) > 1:
            dmax = torch.clamp(abs_delta[valid_mask].max(), min=eps)
            dmin = torch.clamp(abs_delta[valid_mask].min(), min=eps)
            log_dratio = torch.log(dmax) - torch.log(dmin)

            if verbose:
                print(
                    f"[GradRatio] {layer_name:30s} "
                    f"log_ratio_before={log_dratio.item():.4f}"
                )

            if log_dratio > (grad_max_ratio + grad_ratio_tolerance):
                target_dmax = torch.exp(
                    torch.log(dmin) + grad_max_ratio
                )
                clipped_abs_delta = torch.clamp(
                    abs_delta,
                    max=target_dmax,
                )
                delta = torch.sign(delta) * clipped_abs_delta

                if verbose:
                    print(
                        f"[GradRatio] {layer_name:30s} "
                        "large updates clipped"
                    )
            elif verbose:
                print(
                    f"[GradRatio] {layer_name:30s} "
                    "update ratio healthy"
                )


    weight_new = weight_old + delta
    changed = not torch.equal(weight_new, weight_old)

    if changed:
        _copy_matrix_(module, weight_new)
        status["regulated"] = True


    if verbose:
        if log_ratio_before is not None:
            print(
                f"[Spectral] {layer_name:30s} "
                f"log_ratio_before={log_ratio_before:.4f}"
            )
            print(
                f"[Spectral] {layer_name:30s} "
                f"{correction_message}"
            )
            print(
                f"[Spectral] {layer_name:30s} "
                f"log_ratio_after={log_ratio_after:.4f}"
            )

        if mask_active:
            print(
                f"[Mask] {layer_name:34s} "
                f"active={keep_count}/{len(s_target)} "
                f"masked={status['masked_count']} "
                f"gamma={gamma:.1e}"
            )

        if frozen_ratio is not None:
            print(
                f"[Mask] {layer_name:34s} "
                f"frozen_ratio={frozen_ratio:.6f}"
            )

        if status["mask_triggered"]:
            print(
                f"[Mask] {layer_name:34s} MASK TRIGGERED at "
                f"val_loss={float(current_val_loss):.6f} <= "
                f"{float(mask_after_val_loss):.6f}"
            )

        print("-" * 70)

    return status


@torch.no_grad()
def apply_dynamic_spectral_regulation(
    model: nn.Module,
    *,


    max_ratio: float = 2.0,
    min_ratio: float = 1.0,
    ratio_tolerance: float = 0.1,
    eta: float = 1.0,


    gamma: float = 1.0e-6,


    grad_max_ratio: Optional[float] = 2.0,
    grad_ratio_tolerance: float = 0.1,


    current_val_loss: Optional[float] = None,
    mask_after_val_loss: Optional[float] = None,
    mask_threshold: float = 0.1,


    eps: float = 1.0e-8,
    verbose: bool = False,
) -> Dict[str, int]:

    summary = {
        "layers_seen": 0,
        "layers_regulated": 0,
        "mask_triggered_layers": 0,
        "mask_active_layers": 0,
        "masked_singular_values": 0,
        "svd_failures": 0,
    }

    for name, module in model.named_modules():
        if not _is_supported_matrix(module):
            continue

        summary["layers_seen"] += 1

        result = dynamic_band_spectral_regulate(
            module,
            name,
            max_ratio=max_ratio,
            min_ratio=min_ratio,
            ratio_tolerance=ratio_tolerance,
            eta=eta,
            gamma=gamma,
            grad_max_ratio=grad_max_ratio,
            grad_ratio_tolerance=grad_ratio_tolerance,
            current_val_loss=current_val_loss,
            mask_after_val_loss=mask_after_val_loss,
            mask_threshold=mask_threshold,
            eps=eps,
            verbose=verbose,
        )

        summary["layers_regulated"] += int(bool(result["regulated"]))
        summary["mask_triggered_layers"] += int(bool(result["mask_triggered"]))
        summary["mask_active_layers"] += int(bool(result["mask_active"]))
        summary["masked_singular_values"] += int(result["masked_count"])
        summary["svd_failures"] += int(bool(result["svd_failed"]))

    return summary
```
